# Imputing missing RNA-seq data from DNA methylation by using transfer learning based neural network

**DOI:** 10.1101/803692

**Authors:** Xiang Zhou, Hua Chai, Huiying Zhao, Ching-Hsing Luo, Yuedong Yang

## Abstract

**Background:** Gene expression plays a key intermediate role in linking molecular features at DNA level and phenotype. However, due to various limitations in experiments, the RNA-seq data is missing in many samples while there exists high-quality of DNA methylation data. As DNA methylation is an important epigenetic modification to regulate gene expression, it can be used to predict RNA-seq data. For this purpose, many methods have been developed. A common limitation of these methods is that they mainly focus on single cancer dataset, and do not fully utilize information from large pan-cancer dataset.

**Results:** Here, we have developed a novel method to impute missing gene expression data from DNA methylation data through transfer learning-based neural network, namely TDimpute. In the method, the pan-cancer dataset from The Cancer Genome Atlas (TCGA) was utilized for training a general model, which was then fine-tuned on the specific cancer dataset. By testing on 16 cancer datasets, we found that our method significantly outperforms other state-of-the-art methods in imputation accuracy with 7%-11% increase under different missing rates. The imputed gene expression was further proved to be useful for downstream analyses, including the identification of both methylation-driving and prognosis-related genes, clustering analysis, and survival analysis on the TCGA dataset. More importantly, our method was indicated to be useful for general purpose by the independent test on the Wilms tumor dataset from the Therapeutically Applicable Research To Generate Effective Treatments (TARGET) project.

**Conclusions:** TDimpute is an effective method for RNA-seq imputation with limited training samples.

## Background

Recent development of molecular biology and high-throughput technologies facilitates the simultaneous measurement of various biological omics data such as genomics, transcriptomics, epigenetics, proteomics, and metabolomics for a single patient. Compared with single-omics analysis, integrative analyses of multi-omics data provide comprehensive insights of cancer occurrence and progression, and thus strengthen our ability to predict cancer prognosis and to discover various levels of biomarkers. However, due to limitations of experimental techniques and relatively high costs for measuring the multi-omics data, most samples aren’t measured with all types of omics data, and lack one part of omics types (called “block missing”). This problem is prevalent in publicly available multi-omics datasets including The Cancer Genome Atlas (TCGA). Since gene expression affects clinical outcomes or phenotypes more directly than molecular features at DNA level like DNA methylation and genetic variants [1], we focused on the gene expression data imputation from DNA methylation data.

When the missing data happens at random positions in single omics data, the missing data can be imputed by traditional methods, such as singular value decomposition imputation (SVD), k-nearest neighbor (KNN) [2]. However, these methods don’t perform well when the whole RNA-seq data is missing. In order to address this issue, Voillet et al. developed a multiple hot-deck imputation approach to impute missing rows in multi-omics dataset for multiple factor analysis [3]. Imbert et al. used multiple hot-deck imputations to improve the reliability of gene network inference [4]. In these two methods, they measured the similarities to cases in a standard database, and fixed the missing values according to the case with the highest similarity. As the most similar case is easy to be affected by random fluctuations in its neighbors, Dong *et al*. developed the TOBMI method by using a k-nearest neighbor weighted method to impute mRNA-missing samples, where the similarity was measured by similarities of DNA methylation data [5]. Obviously, these methods depend strongly on its available neighbors, and are of limited accuracies due to relatively small sample sizes of specific cancer datasets. More importantly, they can’t capture information from other related cancer datasets. Recently, the least absolute shrinkage and selection operator (LASSO) penalized regression was employed to predict gene expression using genetic variants [6] and DNA methylation [7], respectively. However, these linear methods are still limited to capture the non-linear relations between genomic variables.

In recent years, deep neural network has demonstrated its superiority on modeling complex nonlinear relationships and enjoys scalability and flexibility. For the gene expression imputation or prediction, many deep learning models have also been proposed. Chen *et al.* built a multilayer feedforward neural network to predict the expression of target genes from the expression of ∼1000 landmark genes [8]. With the ability to recover partially corrupted input data, denoising autoencoder (DAE) was used to impute missing values in single-cell RNA-seq data [9-11]. Xie *et al.* constructed a similar deep model to infer gene expression from genotypes of genetic variants [12]. Based on convolutional neural network, Zeng *et al.* used promoter sequences and enhancer-promoter correlations to predict gene expression [13]. One obstacle for these deep learning models to multi-omics datasets is the high dimensionality (>20,000 features) in omics data but a small sample size (<1000). Even the TCGA has only hundreds of samples for each cancer type. Thus, it is hard to train an accurate model with millions of parameters in deep learning architecture.

In such scenarios, transfer learning is usually considered as a promising method, where parameters trained for a task with large amount of data are reused as the initialization parameters for a similar task with limited data [14]. In the computer vision community, a common strategy is to pretrain the convolutional neural network (CNN) with ImageNet [15] and then fine-tune its last few layers or all layers (depend on the size of the target dataset) for the target tasks. This pretraining approach has achieved state-of-the-art results on many tasks including object detection [16], image segmentation [17], image classification [18], action recognition [19]. Yosinski *et al.* pointed out the relationship between network structure and the transferability of features [20]. They showed that the deep features transition from general to specific along the network, and transferring higher layers result in significant drop in performance since the features are more specific to source datasets.

For the omics data analysis of cancers, the transfer learning strategy has been applied to different tasks. Li *et al.* built a pan-cancer Cox model for the prediction of survival time, where eight cancer types were combined to assist the training of target cancer dataset [14]. Yousefi *et al.* utilized samples from uterine corpus endometrial carcinoma and ovarian serous carcinoma to augment the target breast cancer dataset to improve the prediction of clinical outcomes [21]. Hajiramezanali *et al.* learned information from the Head and Neck Squamous Cell Carcinoma cancer to subtype lung cancer [22]. Based on the assumption that different types of cancer may share common mechanisms [23, 24], transfer learning is becoming a useful approach for the prediction of missing data by learning from the data of different cancer types.

In this study, we propose a new method to utilize the transfer learning based neural network for imputing gene expression from DNA methylation data, namely TDimpute. Specifically, we first train a neural network based on the pan-cancer dataset to build a general imputation model for all cancers, which is then transferred to target cancer types (Fig 1). To the best of our knowledge, this is the first study to employ the transfer learning for the imputation of gene expression from DNA methylation. The method was shown to be superior to other methods to recover gene expressions for 16 cancer types (see Table 1) in the TCGA at five different missing rates. Our imputed gene expressions were further proven useful with the identification of methylation-driving genes, prognosis-related genes, clustering analysis, and survival analysis through the validations on the TCGA dataset and independent test of the Wilms tumor dataset from the Therapeutically Applicable Research To Generate Effective Treatments (TARGET) project.

**Table 1.** The TCGA cancer types and their sample sizes sorted by size, with the first 16 types used for test.

| Cancers | Full Name | Dataset size |
| --- | --- | --- |
| BRCA | Breast adenocarcinoma | 867 |
| THCA | Thyroid carcinoma | 562 |
| HNSC | head and neck squamous cell carcinoma | 541 |
| LGG | Brain lower grade glioma | 541 |
| PRAD | Prostate adenocarcinoma | 532 |
| LUAD | Lung adenocarcinoma | 477 |
| SKCM | Skin cutaneous melanoma | 472 |
| BLCA | Bladder urothelial carcinoma | 424 |
| LIHC | Liver hepatocellular carcinoma | 416 |
| LUSC | Lung squamous cell carcinoma | 378 |
| STAD | Skin cutaneous melanoma | 371 |
| KIRC | Kidney renal clear cell carcinoma | 342 |
| CESC | Cervical squamous cell carcinoma and endocervical adenocarcinoma | 308 |
| COAD | Colon carcinoma | 297 |
| KIRP | Kidney renal papillary cell carcinoma | 297 |
| SARC | Sarcoma | 262 |
| UCEC | Uterine corpus endometrial carcinoma | 172 |

**Fig 1.**
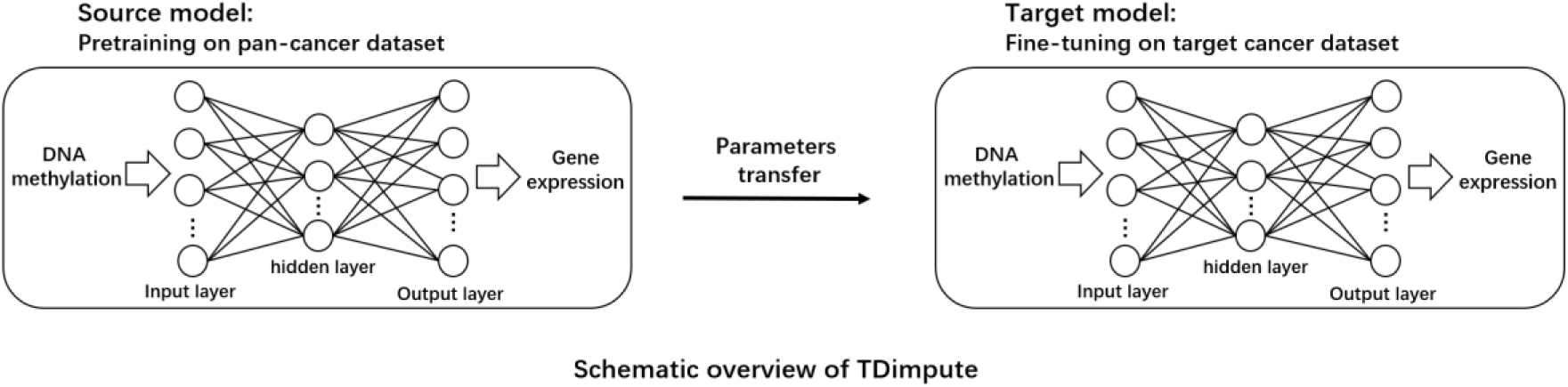
The architecture of transfer learning based neural network (TDimpute) for imputing missing gene expression values in multi-omics dataset. The neural network: DNA methylation data are transformed into gene expression data and the root mean squared error (RMSE) between the actual output and desired output is minimized. Transfer learning: pan-cancer dataset is used to train the general imputation model for pan cancers, which is specifically tuned for each type of cancer.

## Data Description

### Datasets and preprocessing

#### TCGA

We obtained the data of 33 cancer types from The Cancer Genome Atlas (TCGA) using the R package TCGA-assembler [25], including RNA-seq gene expression data (UNC IlluminaHiSeq_RNASeqV2_RSEM), DNA methylation data (JHU-USC HumanMethylation450), and clinical information with follow-up and tumor–node–metastasis (TNM) cancer stages [26]. Originally, 20531 genes and 485577 methylation sites were collected. By excluding genes with zero values in the RNA-seq data across all samples, 19027 genes remained. The gene expressions were converted by the log_2_(G + 1), where G is the raw gene expression value. For DNA methylation data, we excluded methylation sites with “NA” values, and 269023 methylation sites remained. By further removing sites with small variances (< 0.05) over all samples, 27717 CpG sites were kept. Here, for evaluating all imputing methods we kept only samples having both RNA-seq and DNA methylation data. Finally, the dataset contains 8856 samples with expression data for genes and methylation values of 33 cancers, namely TCGA dataset. To keep enough sample size for downstream analyses, we selected cancer types containing > 200 samples with complete DNA methylation, gene expression, and clinical data, leading to 16 cancer types for test (See Table 1).

#### TARGET

Apart from TCGA, we compiled another dataset developed from the Therapeutically Applicable Research To Generate Effective Treatments (TARGET) project. We chose the Wilms tumor (the most common type of childhood kidney cancer) that has the smallest sample size. Here, the DNA methylation data was downloaded from the TARGET Data Matrix [27], and its corresponding RNA-seq data (RSEM estimated read counts) was downloaded from the UCSC Xena [28]. The data was normalized into the same distribution as TCGA through the quantile normalization [29]. Finally, we obtained 118 samples with complete gene expression data, methylation data, and clinical data, which were randomly split into training and test datasets with proportion of 1:1.

## Analyses

### Comparisons on the imputation accuracy

The imputation methods were evaluated by the average values of the root mean square error (RMSE), mean absolute error (MAE), and the squared Pearson correlation coefficient (*R*^2^) across 16 cancer datasets. By selecting one portions (i.e. 1.0 - missing rate) of samples for constructing/fine-tuning the models, the models were then applied to the remained samples. As shown in Fig 2A (Fig S2 for each dataset), Lasso [7] achieved similar but consistently lower RMSE than TOBMI [5], which indicated that penalized regression had better prediction of the regulation between methylation and gene expression. The SVD method [2] performed worse, demonstrating a slow increase of RMSE from 1.06 to 1.10 with the changes of missing rates from 10% to 70%, but then a sharp increase to 1.24 that was even higher than the result by the Mean method. Overall, the Mean method had the worst performance, which coincided with the trend in the previous study [5]. By comparison, TDimpute-self (indicates the TDimpute trained and predicted on the target cancer dataset) without using transfer learning yielded 2%-9% lower RMSE than Lasso at different missing rates. TDimpute-noTF, as a general model, was trained on the pan-cancer dataset (excluding the target cancer). The model didn’t use information from the target cancer and thus showed a constant performance. It didn’t perform well but produced lower imputation RMSE than the Mean method. The RMSE was even lower than that of SVD and TOBMI when the missing rate was above 70%. TDimpute, a further transfer learning on the target cancer from TDimpute-noTF, decreased the RMSE by 7%-16% over TDimpute-noTF. The RMSE by TDimpute was also 2%-5% lower than TDimpute-self with a bigger difference at a higher missing rate. These results confirmed the power of our TDimpute method in transferring knowledge from other cancer types to improve the imputation performances. We also noted SVD, Lasso, and TOBMI had close to constant RMSE values at missing rates from 10% and 70%, indicating that three times of the sample sizes didn’t contribute much to increase in the imputation accuracies. By comparison, deep learning methods, TDimpute and TDimpute-self, decreased the RMSE by 5% and 7%, respectively, indicating the ability of further improvement with an increase of sample sizes in future.

**Fig 2.**
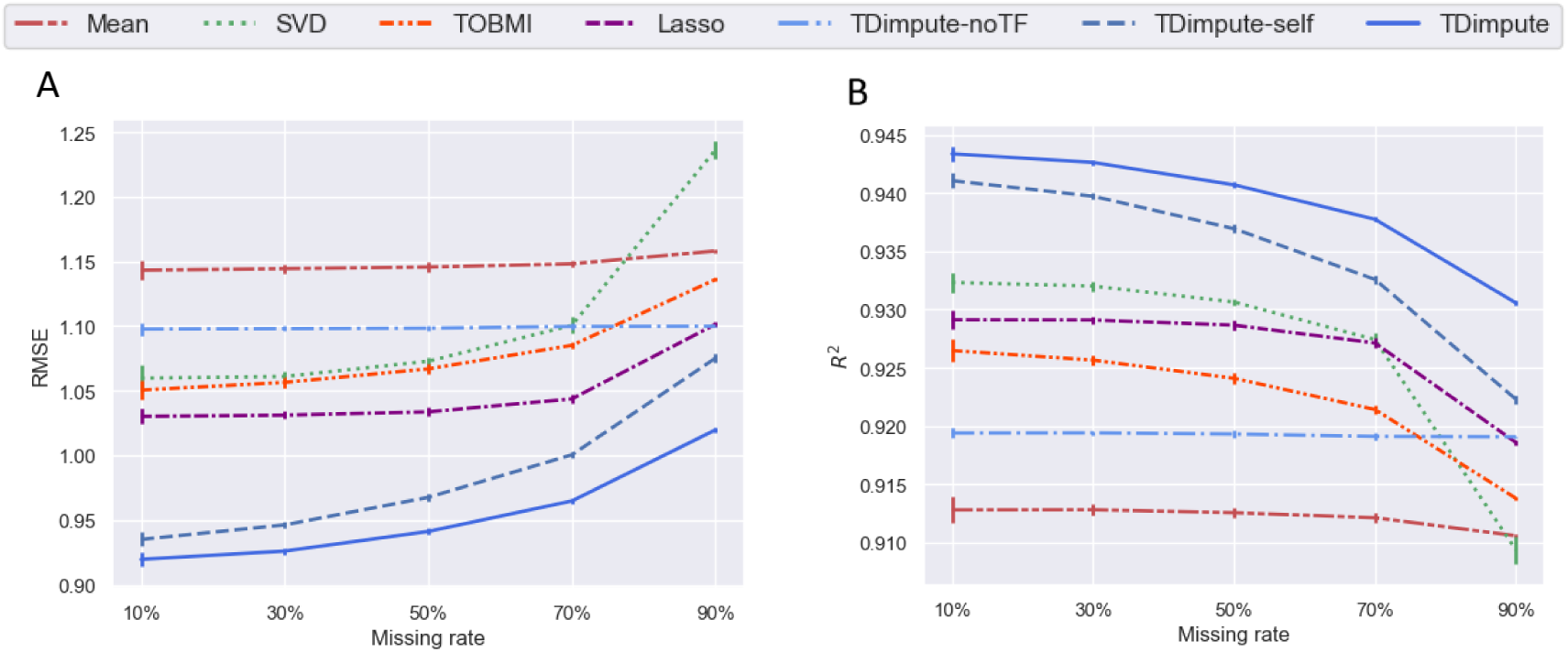
Imputation accuracy of each imputation method. Results were averaged across 16 imputed cancer datasets. (**A)** RMSE values of each method. (**B**) The squared Pearson correlation coefficient (*R*^2^) between each sample of the imputed data and the original full data. TDimpute-self indicates the TDimpute trained and predicted on the target cancer dataset. TDimpute-noTF indicates the TDimpute trained on the pan-cancer dataset (excluding the target cancer) and predicted on the target cancer dataset. The error bar shows the standard deviation.

Since RMSE is sensitive to outliers, we also evaluated the imputation performances with MAE that is less affected by anomalous outliers. As shown in Fig S1, TDimpute consistently performed the best at all missing rates with MAE of 0.616 to 0.685. Slightly differently, TOBMI turned similar to Lasso with lower MAE at 10% while slightly higher MAE at missing rates >50%.

When measured by the squared correlation (*R*^2^) between the imputed and actual values by each sample (Fig 2B, Fig S3 for each dataset), approximatively the same trends could be observed for all methods. Differently, SVD ranked the 3^rd^ except at a missing rate of 90%, where SVD had the lowest *R*^2^ of 0.909. The imputations by the mean gene expressions kept the lowest performance. Hereafter, we will focus on the comparison with SVD, Lasso, and TOBMI methods.

### Impacts on the methylation-expression correlations and the identification of methylation-driving genes

For multi-omics dataset, proper imputation method should preserve the correlation structures between different types of omics. Since the most correlated CpG-gene pairs play the most important roles, we only compared the impact of imputation methods by the average *R*^2^ of top 100 CpG-gene pairs from full datasets. As shown in Fig 3 (Fig S4 for each dataset), all imputation methods cause decreases in the correlations, and differences raise with the missing rates. In general, the TDimpute has the highest recovery power for the methylation-expression correlation with *R*^2^ values close to the actual correlation (*R*^2^=0.68). The performance is followed by TDimpute-self. At a missing rate of 90%, TDimpute-self by using single dataset has a large drop of *R*^2^ to the same level with Lasso. Although TOBMI performs better than SVD according to RMSE, TOBMI consistently has the lowest correlations, likely because TOBMI imputes gene expressions simply by nearest neighbors in DNA methylations that has destroyed the complex methylation-expression relations.

**Fig 3.**
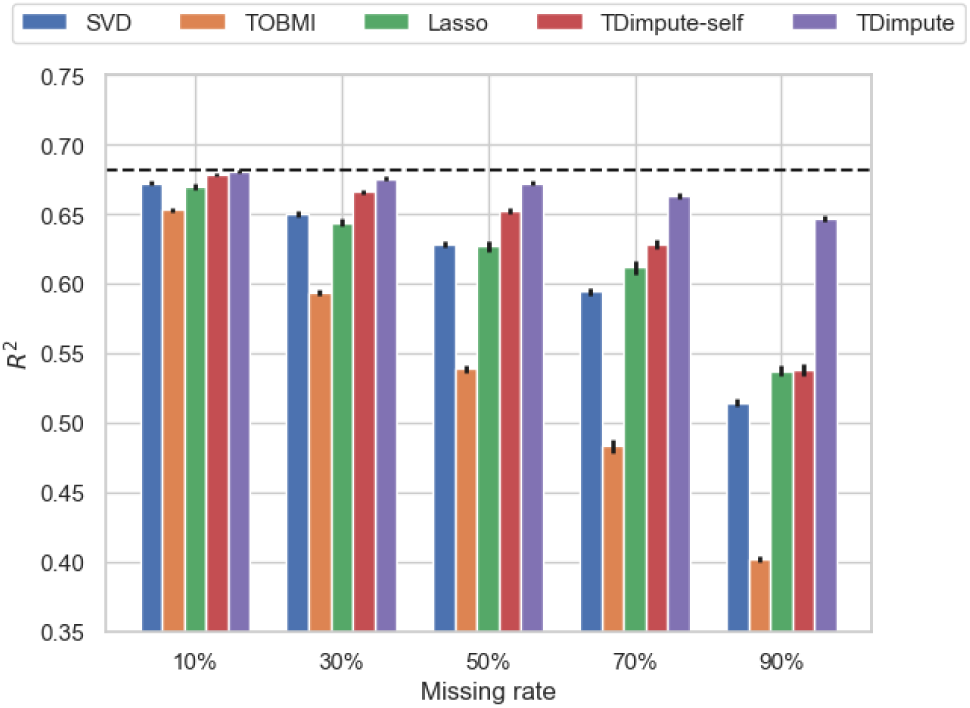
The average correlations *R*^2^ of top 100 CpG-gene pairs over 16 cancer datasets by five imputed methods. Dashed black line is the correlations from the actual dataset. The error bar shows the standard error of the mean.

We further investigated whether the preservation of correlations can obtain better performances in identifications of methylation-driving genes. As shown in Table 2 and Table S2.1, TDimpute consistently shows the superiority in preserving methylation-driving genes from original data with the highest PR-AUC and top 100 overlaps, respectively. TDimpute-self ranks the second in selecting methylation-driving genes, followed by SVD, Lasso, and TOBMI. Compared with TDimpute-self, Tdimpute achieves 0.3%-34% and 1%-49% improvements for PR-AUC and overlap ration, respectively. The improvement is especially pronounced at high missing rates. These results confirm the consistency between correlation preservation and methylation-driving genes identification.

**Table 2.** The average PR-AUC over 16 cancers for recovering methylation-driving genes according to the imputed relative to the actual gene expression data. Average performance across 16 imputed cancer datasets are reported. Best results are highlighted in bold face. The * indicates statistical significance (paired T-test, p-value < 0.05) between TDimpute and other methods.

| Missing rate | SVD | TOBMI | Lasso | TDimpute-self | TDimpute |
| --- | --- | --- | --- | --- | --- |
| 10% | 0.972* | 0.971* | 0.965* | 0.980* | **0.983** |
| 30% | 0.867* | 0.867* | 0.842* | 0.908* | **0.927** |
| 50% | 0.737* | 0.703* | 0.702* | 0.807* | **0.848** |
| 70% | 0.587* | 0.521* | 0.565* | 0.678* | **0.749** |
| 90% | 0.391* | 0.308* | 0.390* | 0.397* | **0.601** |

The PR-AUC and overlap of top 100 methylation-driving genes per cancer dataset are detailed in Table S1 and Table S2.2, respectively.

### Impacts on the identification of prognosis-related genes

We investigated the recovery power of different imputation methods on the identification of significantly prognosis-related genes. To evaluate the selected genes, we compared the PR-AUC and the overlaps between top 100 genes identified from the imputed and the actual data in Table 3 and Table S4.1, respectively. We found that TDimpute consistently achieved the best performances in all missing rates, with 2%-28% higher PR-AUC values and 4%-54% more number of overlapped genes than those by the TOBMI method. Lasso achieved lower values than TOBMI, except at a missing rate of 90%, where Lasso slightly overtook TOBMI. The high agreements and overlap ratios indicate that the imputed gene expressions by TDimpute are more relevant to the clinical outcome.

**Table 3.** The average PR-AUC for recovering prognosis-related genes according to the imputed relative to the actual gene expression data. The * indicates statistical significance (paired T-test, p-value < 0.05) between TDimpute and other methods.

| Missing rate | SVD | TOBMI | Lasso | TDimpute-self | TDimpute |
| --- | --- | --- | --- | --- | --- |
| 10% | 5.53* | 5.83 | 5.55 | 5.71 | **5.91** |
| 30% | 3.46* | 4.08 | 3.71* | 4.22 | **4.25** |
| 50% | 2.08* | 2.74 | 2.42* | 2.94 | **3.06** |
| 70% | 1.14* | 1.60* | 1.33* | 1.87 | **2.03** |
| 90% | 0.56* | 0.47* | 0.49* | 0.89* | **1.14** |

The top 100 genes (ranked by p-values) were additionally compared with the prognosis-related gene list downloaded from The Human Protein Atlas [30] by the enrichment relative to random selections. As shown in Table 4, TDimpute achieved the largest enrichment factors (see Methods section for definition), indicating its ability to identify the really validated prognosis-related genes. At missing rates less than 70%., TOBMI consistently outperformed Lasso and SVD performed the worst. At the missing rate of 90%, all methods except TDimpute performed worse than random selection with enrichment factors less than 1.0.

**Table 4.** The average enrichment factors of top 100 prognosis-related genes overlapped with the genes collected in the Human Protein Atlas. The * indicates statistical significance (paired T-test, p-value < 0.05) between TDimpute and other methods.

| Missing rate | SVD | TOBMI | Lasso | TDimpute-self | TDimpute |
| --- | --- | --- | --- | --- | --- |
| 10% | 0.901* | 0.912* | 0.912* | 0.923* | **0.927** |
| 30% | 0.711* | 0.747* | 0.742* | 0.768* | **0.784** |
| 50% | 0.561* | 0.595* | 0.590* | 0.627* | **0.652** |
| 70% | 0.428* | 0.454* | 0.449* | 0.487* | **0.523** |
| 90% | 0.286* | 0.287* | 0.292* | 0.311* | **0.376** |

The PR-AUC, overlap of top 100 prognostic genes, and the enrichment factors per cancer dataset are detailed in Table S3, S4.2, and S5, respectively.

### Impacts on the performances of clustering analysis and survival analysis

We also evaluated the effects of different imputation methods on clustering analysis and survival analysis. By input of top 100 prognosis-related genes, K-means algorithm was used to divide the samples into two clusters. Fig 4A (Fig S5 for each dataset) shows the adjusted rand index (ARI) for evaluating the concordance between clusters from the imputed and actual gene expression. For all methods, accuracy decreases with the increase of the missing rate, agreeing with the previous study [10]. As expected, TDimpute achieved the highest clustering concordance among the five imputation methods consistently under different missing rates.

**Fig 4.**
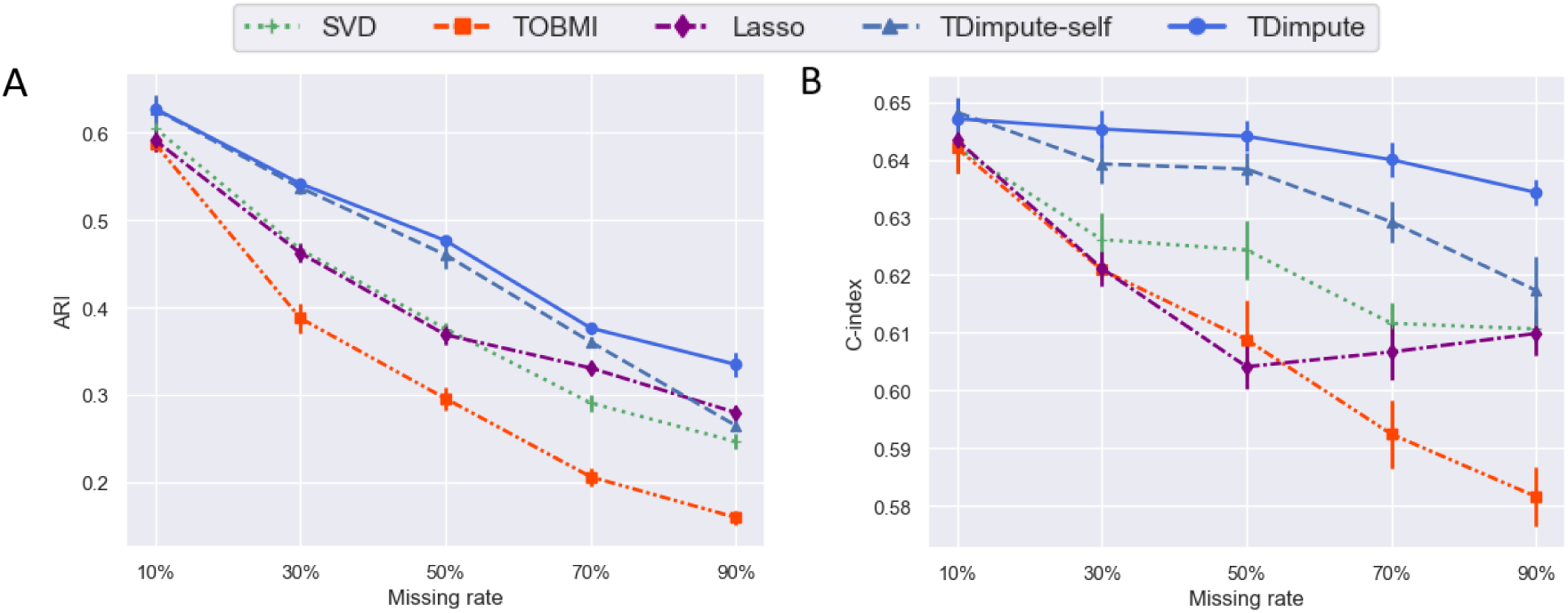
**(A)** The average adjusted rand index (ARI) of the clusters from the imputed and actual data, and **(B)** the average C-index by survival analyses based on imputed data over 16 cancers. The error bar shows the standard error of the mean.

A further survival analysis (Fig 4B, Fig S6 for each dataset) shows that TDimpute consistently outperforms the competing methods. Interestingly, the C-index is not so sensitive to the missing rates. Even at a missing rate of 90%, imputed data caused a decrease of 2-9% in C-index, much smaller than the decreases of 59%-68% in PR-AUC for recovering prognosis-related genes and 45%-74% in ARI for clustering analysis. This is likely because the Cox models were trained with contributions from multiple gene features, and thus bad imputations could be avoided. Similar results were also found in previous studies [31, 32]. For reference, the C-index was 0.669 according to the TNM typing system [26] that were decided by clinicians by tumor phenotypes, indicating the necessity to combine genotype and phenotype for survival analyses in future.

### Validation on the UCEC from TCGA dataset

As a real data application, we performed survival analysis on the Uterine corpus endometrial carcinoma (UCEC) because it has the largest proportion of missing gene expression in the TCGA (172 samples with RNA-seq and 267 without). By using the imputed samples, TDimpute achieved the highest C-index of 0.588, compared to TDimpute-self, SVD, TOBMI, Lasso with C-index of 0.575, 0.55, 0.553, and 0.508, respectively. Fig S7 shows the survival curves of two groups separated by K-means. The two groups for TDimpute show a significant difference (P=1.7e-04) in the survival curves according to the log-rank test. By comparison, the p-values are 1.1e-03, 1.9e-03, 9.5e-03, and 1.2e-03, for TDimpute-self, Lasso, TOBMI, and SVD, respectively.

### Independent test on the TARGET dataset

As an independent test beyond TCGA, we selected the smallest Wilms tumor from TARGET project. By fine-tuning the TCGA pan-cancer model with randomly selected 59 samples (50% of the dataset), the model was tested on the left samples. As expected, TDimpute achieved the lowest RMSE of 0.955 (TDimpute-self: 0.98; SVD: 1.064; Lasso: 1.006; TOBMI: 1.018). This demonstrates the generalization of the TCGA-pretrained model on independent dataset. K-means method is used to cluster the 118 samples after imputation, and two resulted clusters were used to plot the survival curves. For the Kaplan-Meier survival curves in Fig 5, TDimpute achieved the best prognostic stratification with P-value of 7.81e-08, compared to 4.53e-07, 8.64e-06, 3.80e-03, 2.99e-02 for TDimpute-self, TOBMI, Lasso, and SVD, respectively. TDimpute achieved the highest C-index of 0.592, compared to 0.591, 0.523, 0.558, and 0.501 for TDimpute-self, SVD, TOBMI, Lasso, respectively.

**Fig 5.**
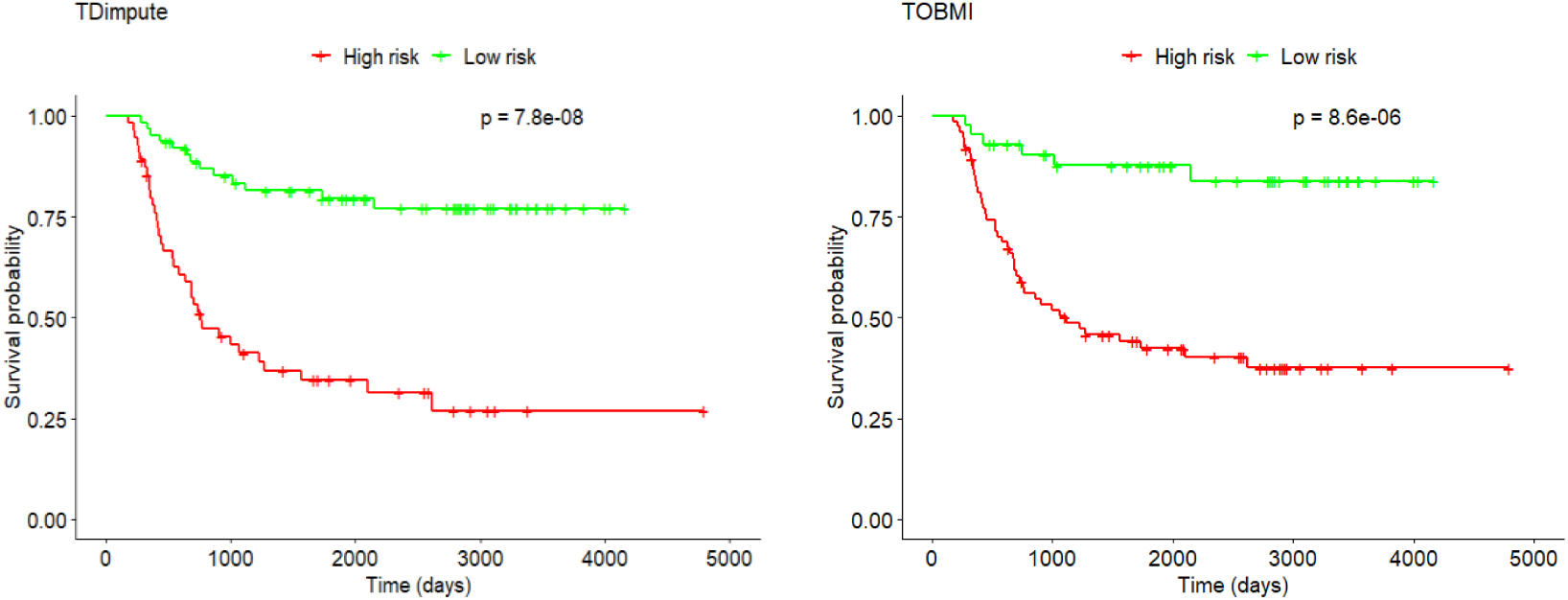
Kaplan-Meier plot for the two clusters obtained from the Wilms tumor dataset imputed by TDimpute and TOBMI, respectively.

## Discussion

In this study, TDimpute performs missing gene expression imputation by building a highly nonlinear mapping from DNA methylation to gene expression data. Due to the limited size of cancer datasets in TCGA, we employed transfer learning to capture the commonalties in pan-cancer dataset for pre-training parameters. We compared TDimpute with/without transfer learning, Lasso, SVD, and TOBMI methods by RMSE, MAE, and correlation *R*^2^, methylation-expression correlations. Since the main task of imputations is to recover biologically meaningful gene expression data for downstream analysis, we also evaluated the impact of imputations on the identification of methylation-driving genes, prognosis-related genes, clustering analysis, and survival prognosis. It is worthy to note that although only methylation and gene expression data are illustrated in this study, our framework is capable of incorporating other omics data.

Experimental results on 16 cancer datasets confirmed that our TDimpute method without transfer learning already outperformed Lasso, SVD, and TOBMI method in different evaluation metrics. By inclusion of transfer learning, the TDimpute method can further improve the performances especially at high missing rates. In addition, the ranking of Lasso, SVD, and TOBMI methods through imputations accuracies (RMSE, MAE, and correlation *R*^2^) are not strictly agreeing with their performances in preservation of methylation-expression correlations, clustering analysis, and survival prognosis, but our TDimpute method performs consistently the best in both the imputation accuracies and downstream analyses.

Besides the good performances of imputation accuracies and downstream analyses, another main advantage of our proposed method is its computational efficiency and convenience. Based on GPU acceleration, our TDimpute method is capable of processing large-scale pan-cancer dataset including tens of thousands of samples and hundreds of thousands of features, while Lasso, TOBMI, and SVD suffer from poor scalability due to the computational complexity of distance matrix computations and singular value decomposition operations. Based on the pre-trained model, transfer learning framework can also accelerate the training process on the target dataset.

In previous study of genome-wide association analysis (GWAS) without directly measured gene expression [6, 33], gene expressions could be imputed from genomic data to perform transcriptome-wide association analysis (TWAS) that can reduce multiple-testing burden and identify associated genes. Recently, many studies have been proposed to impute gene expressions from genomic data, or even from pathology images [34]. In future, the predicted gene expression from other omics data, such as genomics, pathology, or/and radiomics, can also be integrated in epigenome-wide association studies (EWAS) [35].

Future work can focus on reducing the amount of model parameters and integrating more related training samples. Since we only used the correlations between omics for imputations, one possible direction is to leverage prior knowledge of gene-gene interaction network. The known relationships between variables/genes has demonstrated its ability to significantly reduce model parameters by enforcing sparsity on the connections of neural network [36]. The performance of this approach is dependent on the quality of the gene-gene networks, and more investigation need to be done in this direction.

In the preprocessing step, we removed sites with low-variance DNA methylation over all samples, which might remove some sites that are useful for specific cancer types. To test whether the cancer-specific CpG sites from the pan-cancer dataset are necessary for gene expression prediction, we updated the neural networks by taking the top 20000 variable CpG sites specific to each cancer as auxiliary input (Fig S8). Fig S9 shows that the improvement is relatively limited with only 0.4%-1.4% decrease of RMSE, compared to the network without including the cancer-specific sites. These results indicate that the CpG sites with high variances (> 0.05) over all samples are adequate enough for predicting gene expression across different cancers.

## Methods

### Network architecture and model training

#### Neural network architecture

As shown in Fig 1, TDimpute is a three-layer neural network with sizes of [27717, 4000, 19027]. The nodes between layers are fully connected and the sigmoid activation function is adopted. The loss function for training is the root mean squared error (RMSE) which minimizes the difference between the experimentally measured and predicted gene expression value:

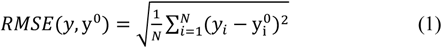

where 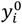 and *y*_*i*_ are the experimentally measured and predicted expression value for gene *i*, and *N* is the dimension of output vector (i.e., the number of genes). The network can be considered as a highly nonlinear regression function that maps DNA methylation data (input) to gene expression data (output).

The network was trained using Adam optimizer with default parameters (learning rate set as 0.0001) [37]. The method was implemented with TensorFlow [38]. All the codes and pretrained pan-cancer models are available on Github: https://github.com/sysu-yanglab/TDimpute.

#### Transfer learning setting

To train the prediction model for one target cancer in the TCGA, the datasets of other cancer types were combined to generate a multi-cancer model that was then fine-tuned by the target cancer data (Fig 1). The data of the target cancer was excluded to train the multi-cancer model as we needed to remove different portions of the data for the target cancer to evaluate our imputation model. During the fine-tune process, we reused the network architecture and all the initialization parameters from the pretrained model. Here, we didn’t freeze any layer, as this demonstrated better performance.

#### Hyper-parameters tuning

For our neural network, we selected BRCA dataset to optimize all hyper-parameters by RMSE through 5-fold cross validation. Here, we optimized the hyper-parameters based on the BRCA dataset and then applied the optimal hyper-parameters to all cancer types. Based on the pan-caner dataset (excluding the BRCA dataset), we first optimized the pretrained model to determine the architecture of neural network, i.e., the number of hidden layers, the hidden layer size, and the epochs to stop training. Details on the performance analysis for each hyper-parameter were provided in Table S6 and Fig S10 in the supplementary material. For the fine-tune process on the BRCA dataset, the only hyper-parameter we need to choose is the training epoch since the network architecture was determined by the pretrained model. Fig S11 shows the convergence process of different levels of missingness on the validation dataset of BRCA.

Finally, we selected the following hyper-parameters for pan-cancer model: 1 hidden layer (selected from 1 and 3) including 4000 nodes (from 500, 1000, 2000, 4000, and 5000), Sigmoid activation function (from Tanh, Relu, and Sigmoid), epochs of 300 (from 50, 100, 150, 300, and 500), and batch size of 128. For the fine-tuning stage, 150 epochs (from 50, 100, 150, 300, and 500) were used and the batch size was set as 16 because of small sample sizes under large missing rates. Dropout wasn’t used as it decreased the performance [39].

### Training and testing datasets

In the transfer learning for each target cancer type of TCGA, 32 cancer types in the TCGA dataset (except the target cancer type) was used as the source domain dataset for pretraining a model. During the test, in order to simulate performances under different missing rates, we used five fractions (10%, 30%, 50%, 70%, 90%). At each missing rate, we randomly selected one fraction (i.e. 1.0 - missing rate) of the target cancer samples to fine-tune the model that were tested on the left samples to predict the gene expression. The predicted values were then compared with the actual values to evaluated the model performances. To remove random fluctuations, we employed the bootstrapping strategy to repeat this process for 5 times and reported the average performances.

### Performance comparison

We compared our method with other imputation methods, including Lasso [7], TOBMI [5], and SVD methods [2]. The default or suggested parameters were used for these methods. We also evaluated the performances by using the mean expression of each gene for reference.

### Preservation of methylation-expression correlations and methylation-driving genes

Here, we used the squared Pearson correlation coefficient *R*^2^ to evaluate the effect of imputation methods on the correlations between DNA methylation and gene expression. For each gene, we only considered the CpG site with the strongest correlation. Based on the methylation-expression regulation, many studies have been conducted to identify cancer-related DNA methylation-driving (hyper and hypo methylated) genes [40]. Hence, we also evaluated the effects of imputation methods on the identification of methylation-driving genes. The methylation-driving genes (i.e., significantly correlated CpG-gene pairs) were defined with the *R*^2^ ≥ 0.5 and FDR-*q* ≤ 0.05. According to the correlated pairs from original gene expression data, we can compute the area under precision-recall curve (PR-AUC). We also computed the overlap between the top 100 ranked genes identified from the imputed datasets and original full datasets.

### Preservation of prognosis-related genes

A common task in the analyses of gene expression data is the identification of prognostic genes. In order to evaluate the effect of different imputation methods on the identification of potentially prognosis-related genes, we built univariate Cox proportional hazard regression models to select statistically significant genes correlated with overall survivals. With the Cox model, each gene is assigned a p-value describing the significance of the relation between the gene and the target cancer, and genes with p-values ≤ 0.05 were defined as prognosis-related genes. Similar to the evaluations by the methylation-driving genes, PR-AUC and overlapped top 100 genes were used to evaluate all imputation methods.

In addition, we compared our identified genes with the list of prognosis-related genes from The Human Protein Atlas (THPA) [30] through the enrichment faction:

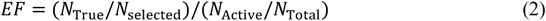

where *N*_True_ is the number of genes appearing in both the THPA and our top *N*_selected_ ranked genes, *N*_Active_ and *N*_Total_ are the number of prognosis-related genes and total number of genes in THPA, respectively.

### Impacts on clustering analysis and survival analysis

We evaluated the relations of genes to cancer survivals by p-values output from the univariate Cox models. By using the top 100 genes, their expression values were used to divide samples into two clusters by the K-means method. The clustering performances were assessed by the adjusted rand index (ARI), a measure of agreement between the predicted clustering labels (by imputed datasets) and the actual clustering labels (by original datasets). We further made survival predictions with significantly related genes (p ≤ 0.05) by using the ridge regression regularized Cox model implemented through the glmnet package [41] in R, a model suitable for fitting regression model with high-dimensional data. The performance of the Cox model was assessed by the Harrell’s concordance index (C-index) that measures the concordance between predicted survival risks and actual survival times.

## Availability of source code and requirements

Project name: Transfer learning for imputing missing RNA-seq data from

DNA methylation

Project home page: https://github.com/sysu-yanglab/TDimpute

Operating system(s): Platform independent

Programming language: Python

License: MIT

biotoolsID: Tdimpute

RRID: SCR_018306

## Availability of supporting data and materials

The data sets and pretrained pan-cancer models supporting the results of this article are available in the Synapse with ID: syn21438134 [42].

## Abbreviations

TCGA: The Cancer Genome Atlas;
TARGET: Therapeutically Applicable Research To Generate Effective Treatments;
SVD: singular value decomposition imputation;
KNN: k-nearest neighbor;
LASSO: least absolute shrinkage and selection operator;
TOBMI: trans-omics block missing data imputation;
DAE: denoising autoencoder;
CNN: convolutional neural network;
TDimpute: transfer learning-based deep neural network for imputation;
RMSE: root mean square error;
MAE: mean absolute error;
ARI: adjusted rand index;
TNM: tumor–node–metastasis;
GWAS: genome-wide association analysis;
EWAS: epigenome-wide association studies;
PR-AUC: area under precision-recall curve;
THPA: The Human Protein Atlas;
C-index: concordance index.

## Consent for publication

Not applicable

## Authors’ Contributions

Y.Y. and C.L. conceived the study. Y.Y. and X.Z. contributed to the design and development of the model. X.Z. and H.C. implemented the software and led the curation of the datasets. Y.Y., H.Z., and X.Z. led the analytics. All authors wrote and approved the final manuscript.

## Acknowledgements

We’d like to acknowledge TCGA and TARGET to make the data publicly available.

## Supplement figures and tables

**Fig S1.**
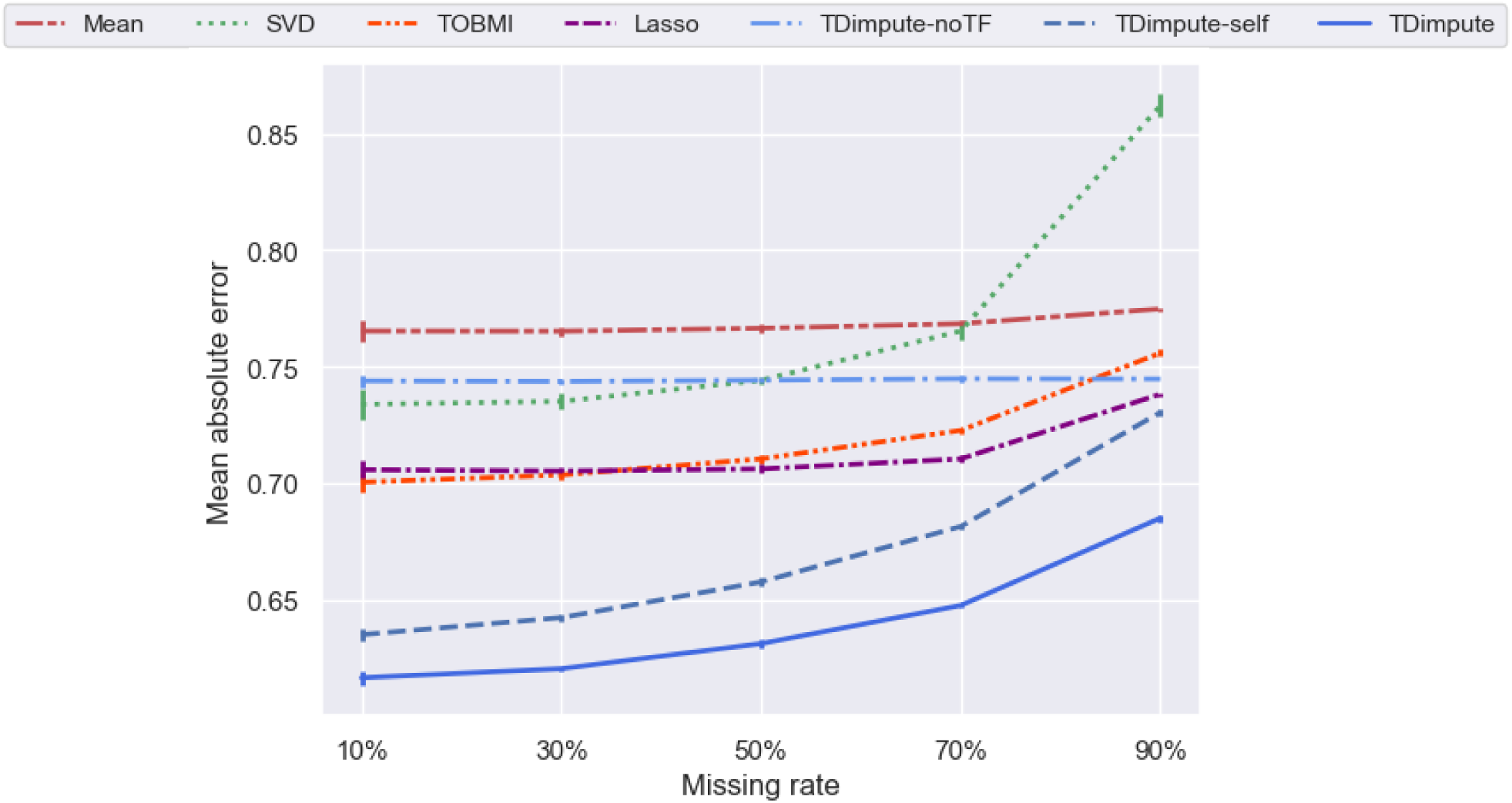
Mean absolute error of each imputation method. Results were averaged across 16 imputed cancer datasets. TDimpute-self indicates the TDimpute trained and predicted on the target cancer dataset. TDimpute-noTF indicates the TDimpute trained on the pan-cancer dataset (excluding the target cancer) and predicted on the target cancer dataset. The error bar shows the standard deviation.

**Fig S2.**
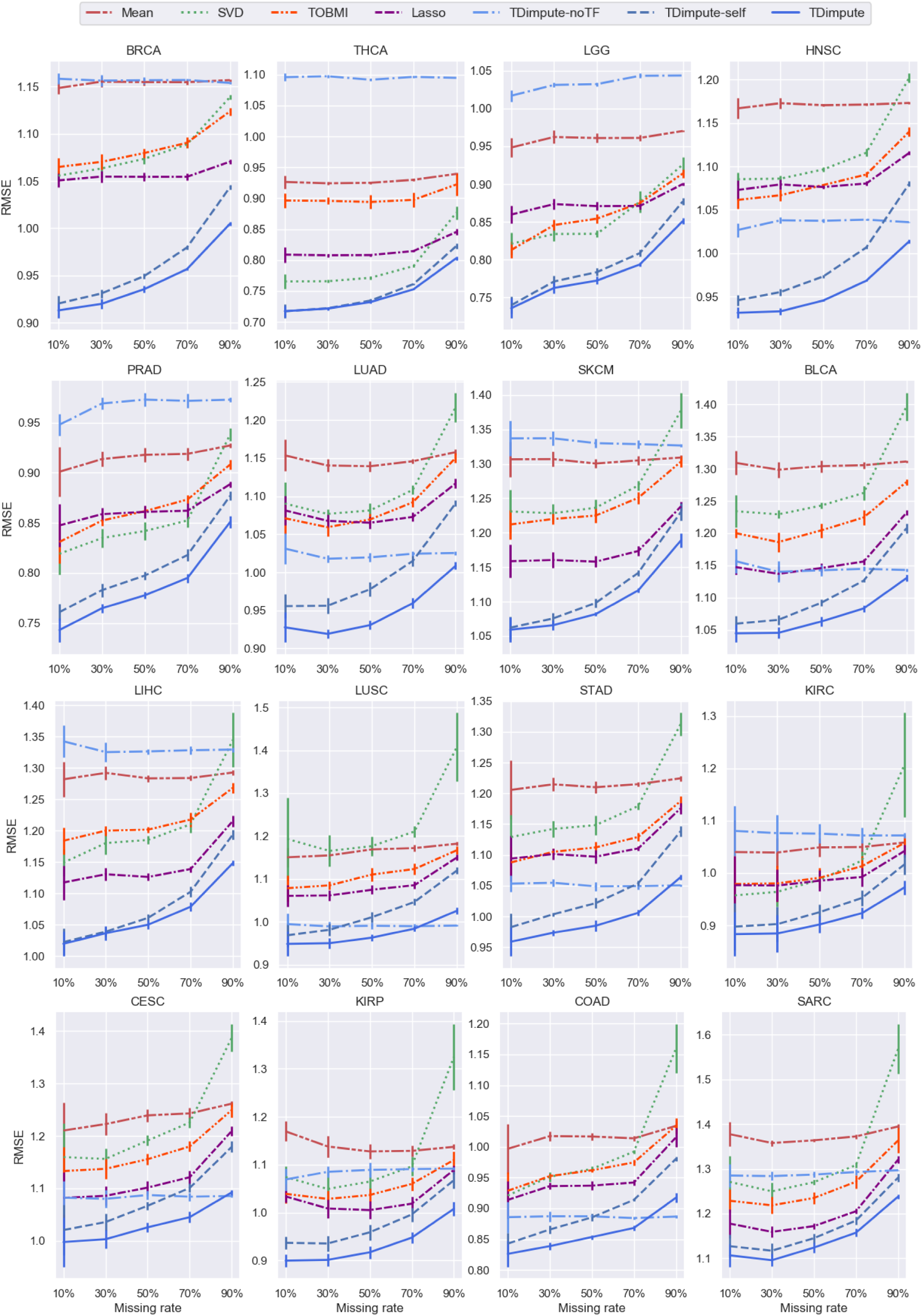
RMSE on 16 imputed cancer datasets with different missing rates. The results were averaged over 5 random replicas. The error bar shows the standard deviation.

**Fig S3.**
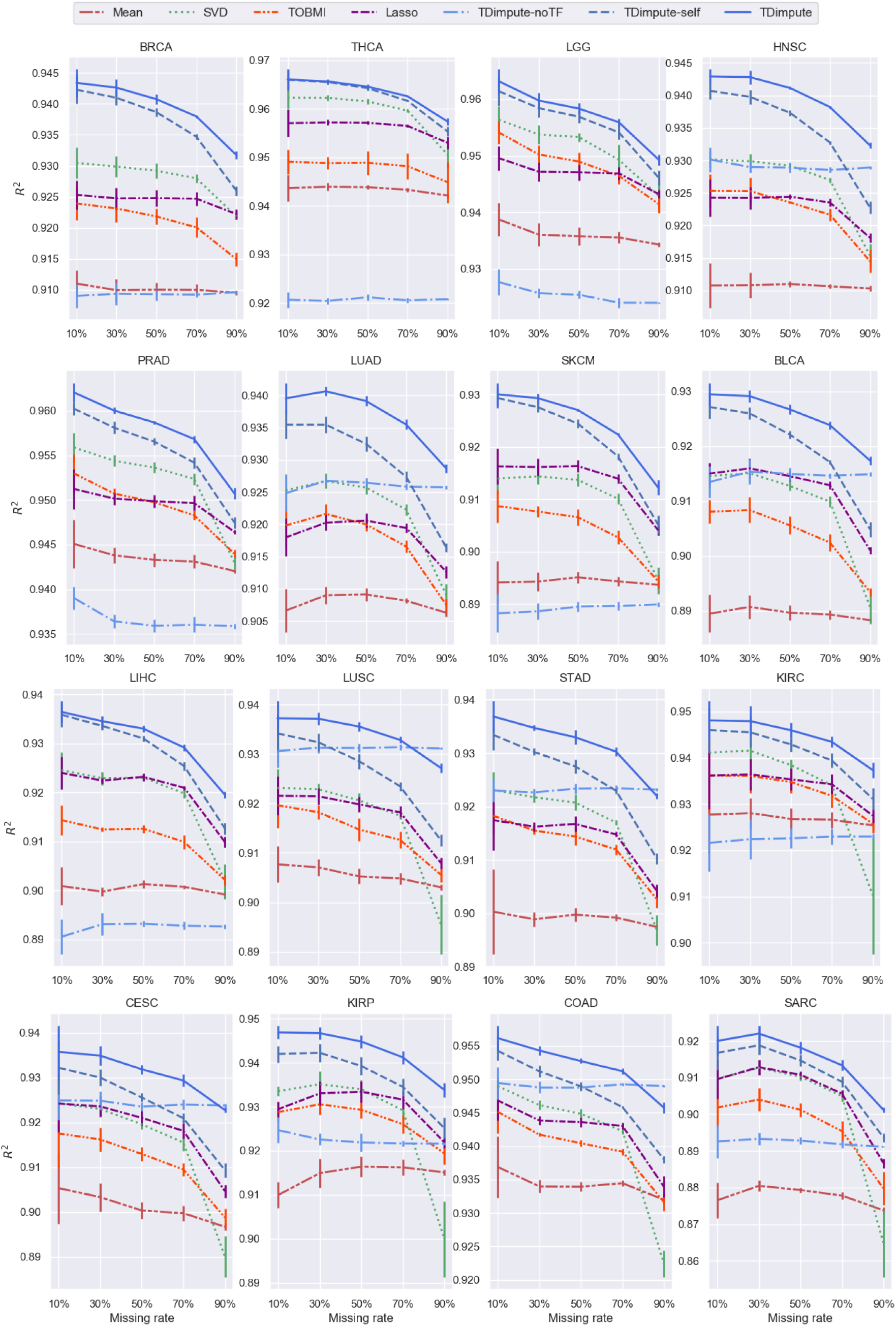
The squared Pearson correlation coefficients *R*^2^ between each sample of the imputed data and the original full data on 16 imputed cancer datasets with different missing rates. The results were averaged over 5 random replicas. The error bar shows the standard deviation.

**Fig S4.**
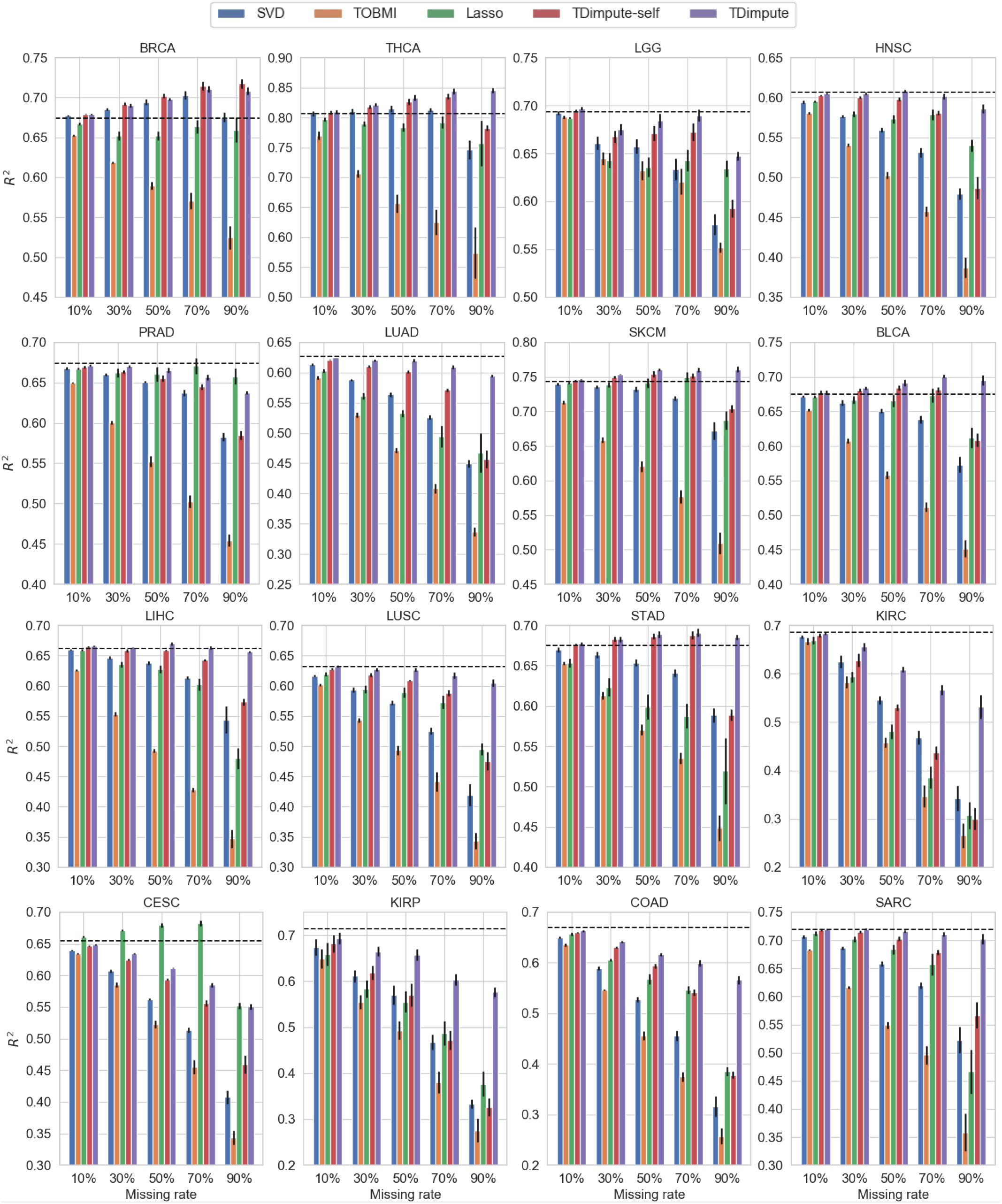
The squared Pearson correlation coefficients *R*^2^ between gene expression and methylation sites on 16 imputed cancer datasets with different missing rates. The results were averaged over 5 random replicas. Dashed black line is drawn as a reference indicating the correlations from the original full dataset. The error bar shows the standard error of the mean.

**Fig S5.**
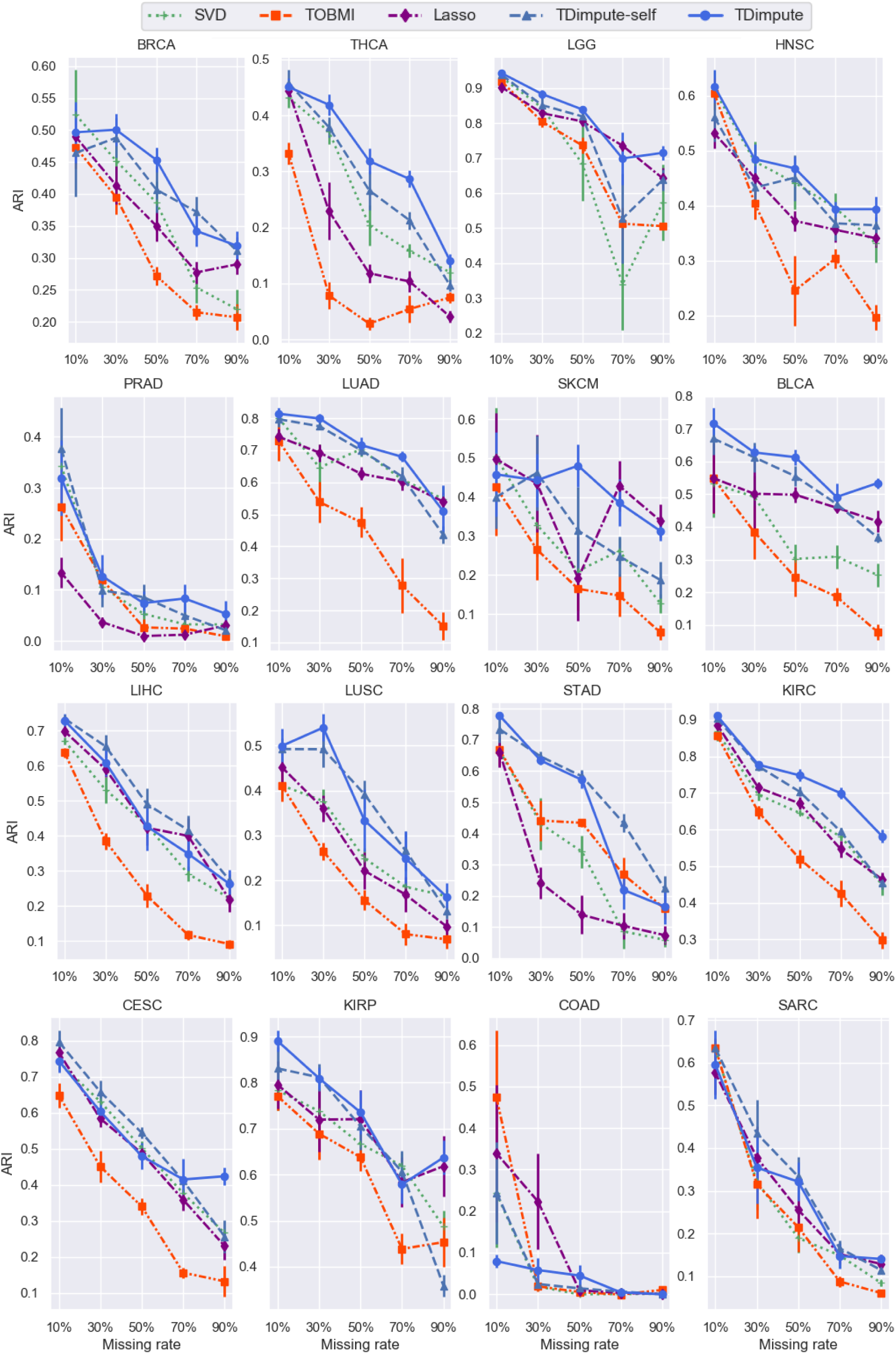
ARI on 16 imputed cancer datasets with different missing rates. The results were averaged over 5 random replicas. The error bar shows the standard error of the mean.

**Fig S6.**
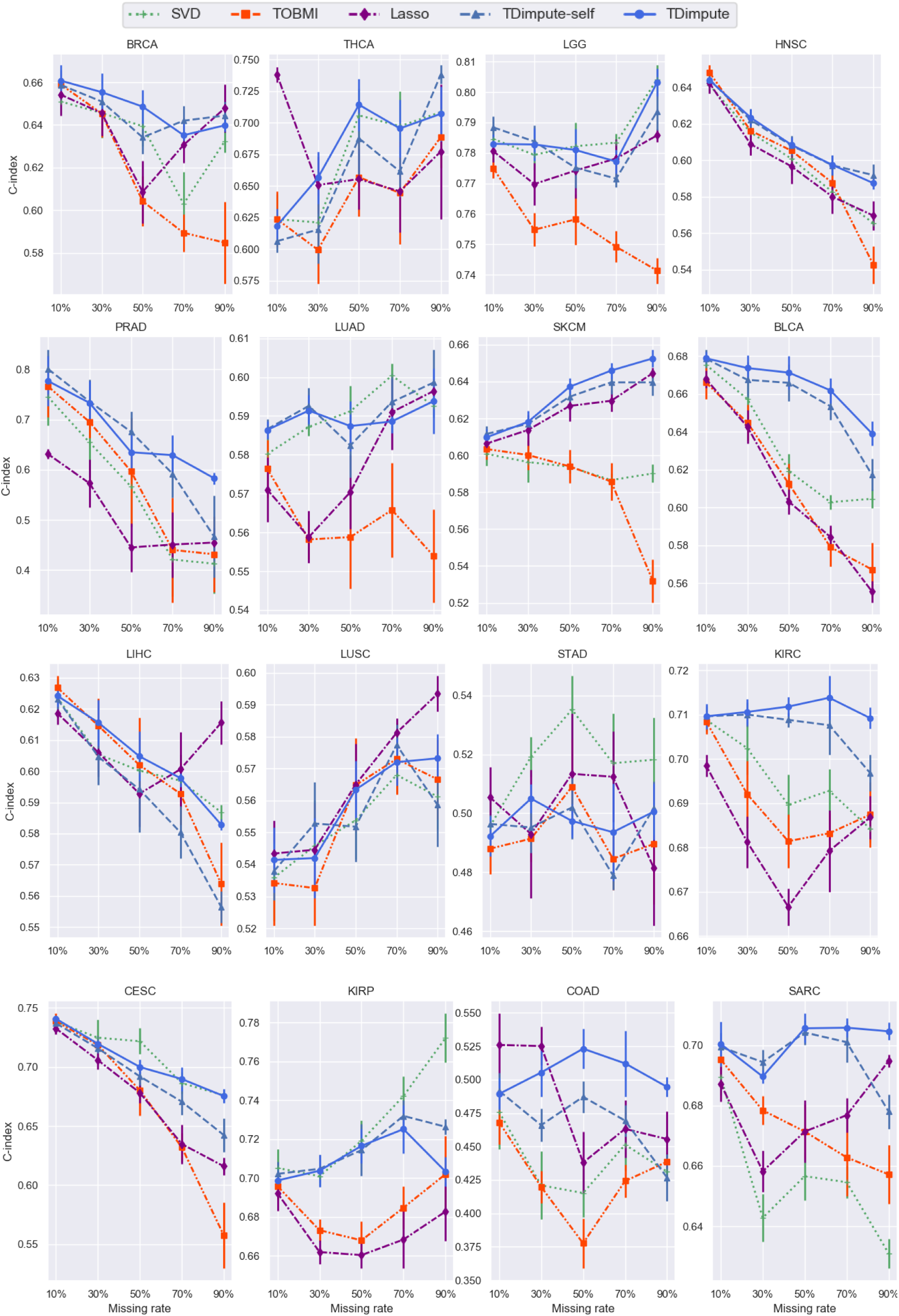
C-index on 16 imputed cancer datasets with different missing rates. The results were averaged over 5 random replicas. The error bar shows the standard error of the mean.

**Table S1.**
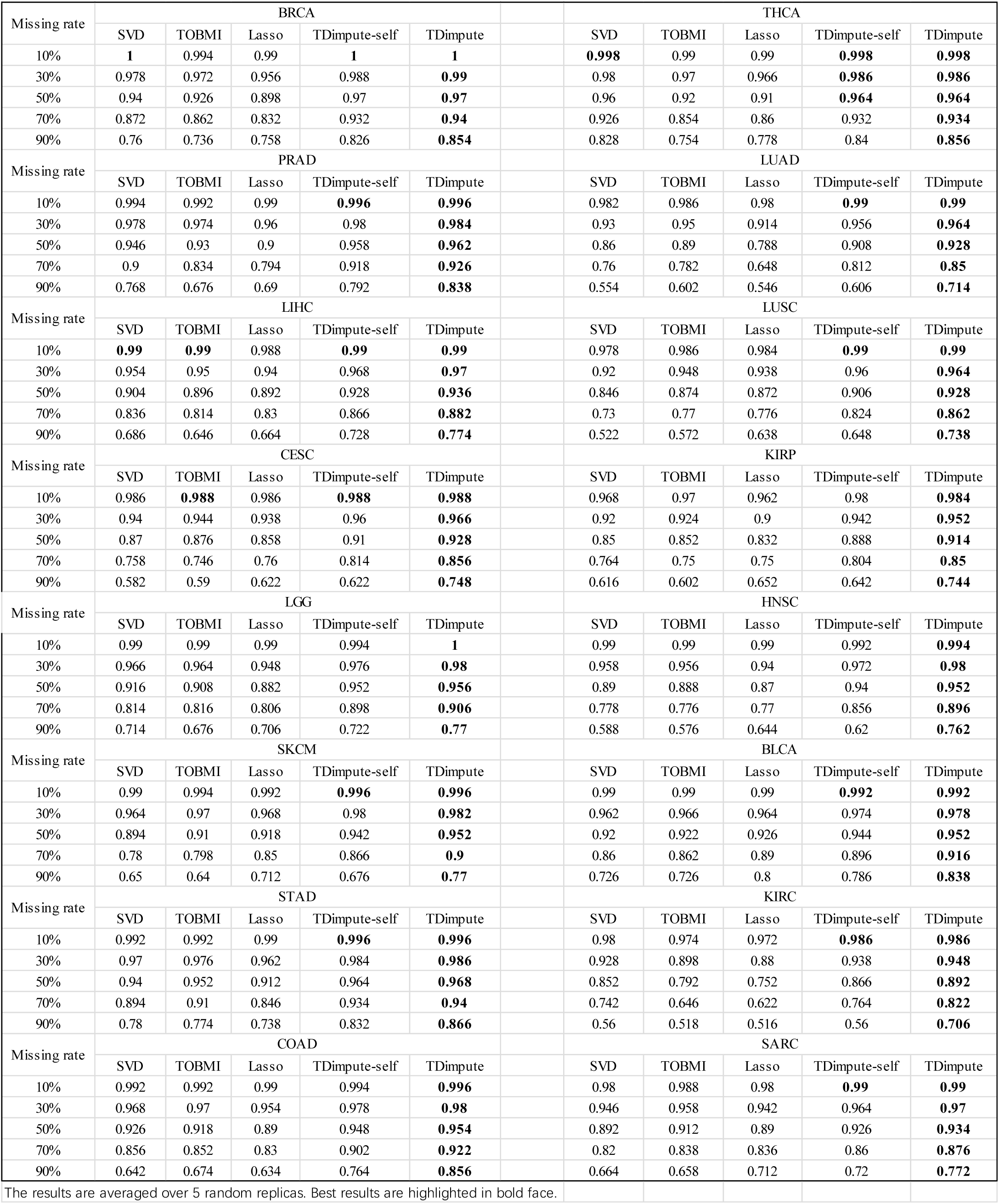
PR-AUC for detecting methylation-driving genes on imputed cancer datasets over 16 cancer types.

**Table S2.1.** Overlap of top 100 methylation-driving genes from imputed dataset and full dataset

| Missing rate | SVD | TOBMI | Lasso | TDimpute-self | TDimpute |
| --- | --- | --- | --- | --- | --- |
| 10% | 89.61* | 87.66* | 86.45* | 91.59 | **91.98** |
| 30% | 76.03* | 71.83* | 68.15* | 83.14* | **84.00** |
| 50% | 62.44* | 55.21* | 52.43* | 73.09* | **76.24** |
| 70% | 45.43* | 37.70* | 37.70* | 57.96* | **65.66** |
| 90% | 24.83 | 18.48* | 22.56* | 26.73* | **50.43** |

The results are averaged over 5 random replicas. Best results are highlighted in bold face.
\* indicates statistical significance (p-value < 0.05) between TDimpute and other methods.

**Table S2.2.** Overlap of top 100 methylation-driving genes between imputed dataset and full dataset over 16 cancer types

| Missing rate | BRCA |  |  |  |  | THCA |  |  |  |  |
| --- | --- | --- | --- | --- | --- | --- | --- | --- | --- | --- |
|  | SVD | TOBMI | Lasso | TDimpute-self | TDimpute | SVD | TOBMI | Lasso | TDimpute-self | TDimpute |
| 10% | 96.2 | 93 | 92.8 | 95.8 | **95.2** | **97.6** | 96 | 94.8 | 97.4 | **97.6** |
| 30% | 93.2 | 82.8 | 80.2 | 92.2 | **93.4** | 94 | 87.2 | 88.8 | **94.8** | 94.2 |
| 50% | 86.8 | 69.4 | 71.8 | **90.4** | 88.8 | 88 | 68.2 | 77.2 | 89.8 | **90.8** |
| 70% | 73.6 | 44 | 53.4 | 81.6 | **85.8** | 75.6 | 57.6 | 66.8 | 83.2 | **85.4** |
| 90% | 34.6 | 9.6 | 21.4 | 45.6 | **66.2** | 34.2 | 31.6 | 41.2 | 52.4 | **71** |
| Missing rate | PRAD |  |  |  |  | LUAD |  |  |  |  |
|  | SVD | TOBMI | Lasso | TDimpute-self | TDimpute | SVD | TOBMI | Lasso | TDimpute-self | TDimpute |
| 10% | 84.4 | 81 | 77.6 | 89 | **90.2** | 93.8 | 92.4 | 77.6 | **94.6** | 93.8 |
| 30% | 61.8 | 59.4 | 54.6 | **78.8** | 77.4 | 84.4 | 82.4 | 54.6 | 87.2 | **87.6** |
| 50% | 48.4 | 47.8 | 42.6 | **66.2** | 64.4 | 77.8 | 73.2 | 42.6 | 82.2 | **83.6** |
| 70% | 38.6 | 37 | 35.2 | 51.2 | **52.6** | 63.8 | 63.4 | 35.2 | 73.8 | **77.8** |
| 90% | 26.2 | 21.8 | 27 | 26.2 | **36** | 40.2 | 38.6 | 27 | 38.6 | **68.4** |
| Missing rate | LIHC |  |  |  |  | LUSC |  |  |  |  |
|  | SVD | TOBMI | Lasso | TDimpute-self | TDimpute | SVD | TOBMI | Lasso | TDimpute-self | TDimpute |
| 10% | 85.6 | 87.2 | 91.4 | 91.4 | **92** | 94.6 | 93.2 | 93.6 | 94 | **95.4** |
| 30% | 72.6 | 72.4 | 68.8 | 80.6 | **81.6** | 86.4 | 83.4 | 75 | **89.6** | 89 |
| 50% | 64.8 | 53.2 | 50 | 72.4 | **76.2** | 75.6 | 71 | 55.8 | 82.6 | **83.8** |
| 70% | 50 | 25.2 | 24.6 | 56.6 | **60** | 51.4 | 46.2 | 32 | 75.2 | **78.4** |
| 90% | 27.4 | 8.2 | 11.6 | 31 | **51.2** | 18.2 | 16.2 | 13.6 | 23.2 | **65** |
| Missing rate | CESC |  |  |  |  | KIRP |  |  |  |  |
|  | SVD | TOBMI | Lasso | TDimpute-self | TDimpute | SVD | TOBMI | Lasso | TDimpute-self | TDimpute |
| 10% | 89.2 | 82.4 | 81.2 | **91** | 90.2 | 83.2 | 79.4 | 78.6 | 84.8 | **87.2** |
| 30% | 73.2 | 64.2 | 60.8 | 78.8 | **82.2** | 68.4 | 57.8 | 60.2 | 74 | **78.2** |
| 50% | 47 | 40.8 | 44.6 | 57.6 | **70.4** | 53.2 | 45 | 50.2 | 59.6 | **67** |
| 70% | 30.4 | 30.6 | 35.6 | 33.2 | **54.2** | 37.6 | 22.8 | 29.4 | 42.8 | **54** |
| 90% | 18.2 | 16.4 | 20.4 | 17.8 | **38.4** | 16.2 | 10.6 | 17.2 | 12 | **36.6** |
| Missing rate | LGG |  |  |  |  | HNSC |  |  |  |  |
|  | SVD | TOBMI | Lasso | TDimpute-self | TDimpute | SVD | TOBMI | Lasso | TDimpute-self | TDimpute |
| 10% | 81.6 | 81.8 | 77.2 | 88.4 | **89.8** | 91.6 | 90.6 | 86.8 | **92.2** | 92 |
| 30% | 67.2 | 63.6 | 56.8 | 75.8 | **79.2** | 80 | 79.8 | 66.4 | **84.6** | 83.4 |
| 50% | 59.8 | 57.4 | 52.6 | 64 | **67.6** | 66.2 | 66.6 | 44.8 | 77.6 | **80.4** |
| 70% | 48.2 | 45.2 | 48.6 | 52.2 | **53** | 42.6 | 52.6 | 31.2 | 65.6 | **72.8** |
| 90% | 33.2 | 21.4 | 38 | 26.4 | **34.6** | 18.2 | 30.6 | 16.4 | 22.2 | **58.2** |
| Missing rate | SKCM |  |  |  |  | BLCA |  |  |  |  |
|  | SVD | TOBMI | Lasso | TDimpute-self | TDimpute | SVD | TOBMI | Lasso | TDimpute-self | TDimpute |
| 10% | 95.6 | 94.8 | 95.8 | **96.4** | **96.4** | 94.4 | 92.2 | 92 | **95** | 94.6 |
| 30% | 86.2 | 83.6 | 88.6 | 91.4 | **92.6** | 85.6 | 78.6 | 75.4 | **90.2** | 90 |
| 50% | 78.4 | 72 | 81.8 | 86.4 | **89.6** | 72 | 55.8 | 54.8 | 85.6 | **86.8** |
| 70% | 60.4 | 47.2 | 68.2 | 71 | **78.4** | 51 | 36.2 | 37.2 | 72.2 | **80** |
| 90% | 36 | 24.4 | 44.2 | 43.8 | **64.2** | 33.4 | 23.8 | 18 | 29.8 | **67** |
| Missing rate | STAD |  |  |  |  | KIRC |  |  |  |  |
|  | SVD | TOBMI | Lasso | TDimpute-self | TDimpute | SVD | TOBMI | Lasso | TDimpute-self | TDimpute |
| 10% | 84.4 | 84.2 | 77.2 | **87.4** | **87.4** | 88.4 | 85.2 | 81 | 89.6 | **91** |
| 30% | 69.4 | 69.8 | 60.8 | **81** | 79 | 71.4 | 66.6 | 56.6 | 74.8 | **77.8** |
| 50% | 54.4 | 55.6 | 47.8 | **69.4** | 68.8 | 55.4 | 47.4 | 37.8 | 60.8 | **65.6** |
| 70% | 36.4 | 41.6 | 37.8 | 55.2 | **61.6** | 36.8 | 30.8 | 25.6 | 45.4 | **54** |
| 90% | 19.4 | 15.2 | 32 | 25.8 | **44** | 24.2 | 19.8 | 20.4 | 13.2 | **39.2** |
| Missing rate | COAD |  |  |  |  | SARC |  |  |  |  |
|  | SVD | TOBMI | Lasso | TDimpute-self | TDimpute | SVD | TOBMI | Lasso | TDimpute-self | TDimpute |
| 10% | 83.4 | 81.4 | 82.6 | **86.8** | 86.4 | 89.8 | 87.8 | 91.2 | 91.6 | **92.4** |
| 30% | 47.6 | 55 | 49.2 | 71.4 | **72** | 75 | 62.6 | 79.4 | 85 | **86.4** |
| 50% | 19 | 31.2 | 23.8 | 54.8 | **60.4** | 52.2 | 28.8 | 58.6 | 70 | **75.6** |
| 70% | 5.4 | 12.6 | 10 | 27.4 | **42.6** | 25 | 10.2 | 36.8 | 40.8 | **60** |
| 90% | 3.4 | 2 | 3.2 | 1.2 | **23.8** | 14.2 | 5.4 | 6 | 18.4 | **43** |

The results are averaged over 5 random replicas. Best results are highlighted in bold face.

**Table S3.**
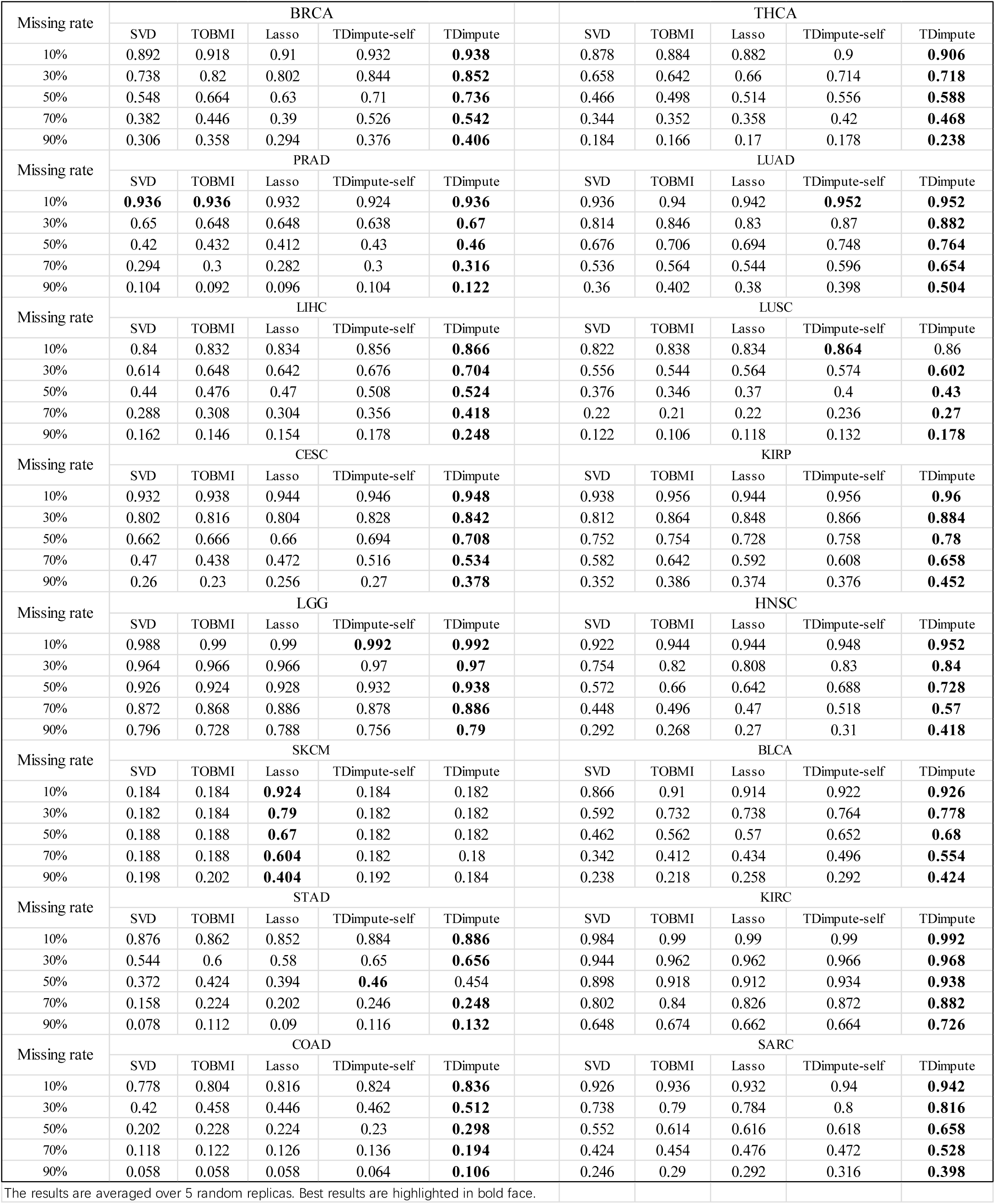
PR-AUC for detecting significantly prognostic gene on imputed datasets over 16 cancer types.

| Missing rate | BRCA |  |  |  |  | THCA |  |  |  |  |
| --- | --- | --- | --- | --- | --- | --- | --- | --- | --- | --- |
|  | SVD | TOBMI | Lasso | TDimpute-self | TDimpute | SVD | TOBMI | Lasso | TDimpute-self | TDimpute |
| 10% | 0.892 | 0.918 | 0.91 | 0.932 | **0.938** | 0.878 | 0.884 | 0.882 | 0.9 | **0.906** |
| 30% | 0.738 | 0.82 | 0.802 | 0.844 | **0.852** | 0.658 | 0.642 | 0.66 | 0.714 | **0.718** |
| 50% | 0.548 | 0.664 | 0.63 | 0.71 | **0.736** | 0.466 | 0.498 | 0.514 | 0.556 | **0.588** |
| 70% | 0.382 | 0.446 | 0.39 | 0.526 | **0.542** | 0.344 | 0.352 | 0.358 | 0.42 | **0.468** |
| 90% | 0.306 | 0.358 | 0.294 | 0.376 | **0.406** | 0.184 | 0.166 | 0.17 | 0.178 | **0.238** |
| Missing rate | PRAD |  |  |  |  | LUAD |  |  |  |  |
|  | SVD | TOBMI | Lasso | TDimpute-self | TDimpute | SVD | TOBMI | Lasso | TDimpute-self | TDimpute |
| 10% | **0.936** | **0.936** | 0.932 | 0.924 | **0.936** | 0.936 | 0.94 | 0.942 | **0.952** | **0.952** |
| 30% | 0.65 | 0.648 | 0.648 | 0.638 | **0.67** | 0.814 | 0.846 | 0.83 | 0.87 | **0.882** |
| 50% | 0.42 | 0.432 | 0.412 | 0.43 | **0.46** | 0.676 | 0.706 | 0.694 | 0.748 | **0.764** |
| 70% | 0.294 | 0.3 | 0.282 | 0.3 | **0.316** | 0.536 | 0.564 | 0.544 | 0.596 | **0.654** |
| 90% | 0.104 | 0.092 | 0.096 | 0.104 | **0.122** | 0.36 | 0.402 | 0.38 | 0.398 | **0.504** |
| Missing rate | LIHC |  |  |  |  | LUSC |  |  |  |  |
|  | SVD | TOBMI | Lasso | TDimpute-self | TDimpute | SVD | TOBMI | Lasso | TDimpute-self | TDimpute |
| 10% | 0.84 | 0.832 | 0.834 | 0.856 | **0.866** | 0.822 | 0.838 | 0.834 | **0.864** | 0.86 |
| 30% | 0.614 | 0.648 | 0.642 | 0.676 | **0.704** | 0.556 | 0.544 | 0.564 | 0.574 | **0.602** |
| 50% | 0.44 | 0.476 | 0.47 | 0.508 | **0.524** | 0.376 | 0.346 | 0.37 | 0.4 | **0.43** |
| 70% | 0.288 | 0.308 | 0.304 | 0.356 | **0.418** | 0.22 | 0.21 | 0.22 | 0.236 | **0.27** |
| 90% | 0.162 | 0.146 | 0.154 | 0.178 | **0.248** | 0.122 | 0.106 | 0.118 | 0.132 | **0.178** |
| Missing rate | CESC |  |  |  |  | KIRP |  |  |  |  |
|  | SVD | TOBMI | Lasso | TDimpute-self | TDimpute | SVD | TOBMI | Lasso | TDimpute-self | TDimpute |
| 10% | 0.932 | 0.938 | 0.944 | 0.946 | **0.948** | 0.938 | 0.956 | 0.944 | 0.956 | **0.96** |
| 30% | 0.802 | 0.816 | 0.804 | 0.828 | **0.842** | 0.812 | 0.864 | 0.848 | 0.866 | **0.884** |
| 50% | 0.662 | 0.666 | 0.66 | 0.694 | **0.708** | 0.752 | 0.754 | 0.728 | 0.758 | **0.78** |
| 70% | 0.47 | 0.438 | 0.472 | 0.516 | **0.534** | 0.582 | 0.642 | 0.592 | 0.608 | **0.658** |
| 90% | 0.26 | 0.23 | 0.256 | 0.27 | **0.378** | 0.352 | 0.386 | 0.374 | 0.376 | **0.452** |
| Missing rate | LGG |  |  |  |  | HNSC |  |  |  |  |
|  | SVD | TOBMI | Lasso | TDimpute-self | TDimpute | SVD | TOBMI | Lasso | TDimpute-self | TDimpute |
| 10% | 0.988 | 0.99 | 0.99 | **0.992** | **0.992** | 0.922 | 0.944 | 0.944 | 0.948 | **0.952** |
| 30% | 0.964 | 0.966 | 0.966 | 0.97 | **0.97** | 0.754 | 0.82 | 0.808 | 0.83 | **0.84** |
| 50% | 0.926 | 0.924 | 0.928 | 0.932 | **0.938** | 0.572 | 0.66 | 0.642 | 0.688 | **0.728** |
| 70% | 0.872 | 0.868 | 0.886 | 0.878 | **0.886** | 0.448 | 0.496 | 0.47 | 0.518 | **0.57** |
| 90% | 0.796 | 0.728 | 0.788 | 0.756 | **0.79** | 0.292 | 0.268 | 0.27 | 0.31 | **0.418** |
| Missing rate | SKCM |  |  |  |  | BLCA |  |  |  |  |
|  | SVD | TOBMI | Lasso | TDimpute-self | TDimpute | SVD | TOBMI | Lasso | TDimpute-self | TDimpute |
| 10% | 0.184 | 0.184 | **0.924** | 0.184 | 0.182 | 0.866 | 0.91 | 0.914 | 0.922 | **0.926** |
| 30% | 0.182 | 0.184 | **0.79** | 0.182 | 0.182 | 0.592 | 0.732 | 0.738 | 0.764 | **0.778** |
| 50% | 0.188 | 0.188 | **0.67** | 0.182 | 0.182 | 0.462 | 0.562 | 0.57 | 0.652 | **0.68** |
| 70% | 0.188 | 0.188 | **0.604** | 0.182 | 0.18 | 0.342 | 0.412 | 0.434 | 0.496 | **0.554** |
| 90% | 0.198 | 0.202 | **0.404** | 0.192 | 0.184 | 0.238 | 0.218 | 0.258 | 0.292 | **0.424** |
| Missing rate | STAD |  |  |  |  | KIRC |  |  |  |  |
|  | SVD | TOBMI | Lasso | TDimpute-self | TDimpute | SVD | TOBMI | Lasso | TDimpute-self | TDimpute |
| 10% | 0.876 | 0.862 | 0.852 | 0.884 | **0.886** | 0.984 | 0.99 | 0.99 | 0.99 | **0.992** |
| 30% | 0.544 | 0.6 | 0.58 | 0.65 | **0.656** | 0.944 | 0.962 | 0.962 | 0.966 | **0.968** |
| 50% | 0.372 | 0.424 | 0.394 | **0.46** | 0.454 | 0.898 | 0.918 | 0.912 | 0.934 | **0.938** |
| 70% | 0.158 | 0.224 | 0.202 | 0.246 | **0.248** | 0.802 | 0.84 | 0.826 | 0.872 | **0.882** |
| 90% | 0.078 | 0.112 | 0.09 | 0.116 | **0.132** | 0.648 | 0.674 | 0.662 | 0.664 | **0.726** |
| Missing rate | COAD |  |  |  |  | SARC |  |  |  |  |
|  | SVD | TOBMI | Lasso | TDimpute-self | TDimpute | SVD | TOBMI | Lasso | TDimpute-self | TDimpute |
| 10% | 0.778 | 0.804 | 0.816 | 0.824 | **0.836** | 0.926 | 0.936 | 0.932 | 0.94 | **0.942** |
| 30% | 0.42 | 0.458 | 0.446 | 0.462 | **0.512** | 0.738 | 0.79 | 0.784 | 0.8 | **0.816** |
| 50% | 0.202 | 0.228 | 0.224 | 0.23 | **0.298** | 0.552 | 0.614 | 0.616 | 0.618 | **0.658** |
| 70% | 0.118 | 0.122 | 0.126 | 0.136 | **0.194** | 0.424 | 0.454 | 0.476 | 0.472 | **0.528** |
| 90% | 0.058 | 0.058 | 0.058 | 0.064 | **0.106** | 0.246 | 0.29 | 0.292 | 0.316 | **0.398** |

The results are averaged over 5 random replicas. Best results are highlighted in bold face.

**Table S4.1.** Overlap of top 100 significantly prognostic genes identified by univariate Cox model between imputed datasets and full datasets.

| Missing rate | SVD | TOBMI | Lasso | TDimpute-self | TDimpute |
| --- | --- | --- | --- | --- | --- |
| 10% | 73.5* | 74.8* | 68.69* | 76.4* | **77.5** |
| 30% | 50.6* | 52.3* | 46.24* | 56.6 | **57.3** |
| 50% | 37.6* | 37.8* | 31.43* | 44.2 | **44.9** |
| 70% | 27.3* | 26.6* | 21.04* | 33.2* | **35.2** |
| 90% | 15.8* | 16* | 11.59* | 20.6* | **24.6** |

The results are averaged over 5 random replicas. Best results are highlighted in bold face.
\* indicates statistical significance (p-value < 0.05) between TDimpute and other methods.

**Table S4.2.** Overlap of top 100 prognostic genes identified by univariate Cox model between imputed dataset and full dataset over 16 cancer types.

| Missing rate | BRCA |  |  |  |  | THCA |  |  |  |  |
| --- | --- | --- | --- | --- | --- | --- | --- | --- | --- | --- |
|  | SVD | TOBMI | Lasso | TDimpute-self | TDimpute | SVD | TOBMI | Lasso | TDimpute-self | TDimpute |
| 10% | 63.4 | 70 | 67 | **73** | 72.4 | 69.2 | 68.2 | 71.2 | 70.2 | **70.6** |
| 30% | 40.6 | 52.2 | 49.4 | 52.8 | **54.8** | 46 | 42.2 | 44.2 | **49.4** | 49 |
| 50% | 27.2 | 33.8 | 28.4 | 36.2 | **39.4** | 31.2 | 28.2 | 29.4 | 33.4 | **35.2** |
| 70% | 13.8 | 18 | 13 | 21.2 | **24** | 22.2 | 18.4 | 18.4 | **25.2** | 23.8 |
| 90% | 6.2 | 5.8 | 4.8 | 9.4 | **12.6** | **16.4** | 8 | 7.2 | 12.8 | 15.4 |
| Missing rate | PRAD |  |  |  |  | LUAD |  |  |  |  |
|  | SVD | TOBMI | Lasso | TDimpute-self | TDimpute | SVD | TOBMI | Lasso | TDimpute-self | TDimpute |
| 10% | 86 | 85.4 | 85.6 | 83.4 | **85.6** | 75.8 | 76.6 | 75 | 79.8 | **82.8** |
| 30% | 51.4 | 49.2 | 50.6 | 52 | **54.6** | 58 | 53.8 | 53.4 | **66** | 64 |
| 50% | 29.4 | 29 | 29.2 | 32.6 | **33.4** | 48 | 41.4 | 40.4 | 51.8 | **53** |
| 70% | 18.6 | 19 | 16 | 19 | **19.4** | 33.2 | 27 | 26.4 | 37.8 | **47.2** |
| 90% | 2.8 | 3.4 | 3.4 | 3.6 | **4.4** | 19.4 | 9.8 | 14 | 21 | **32** |
| Missing rate | LIHC |  |  |  |  | LUSC |  |  |  |  |
|  | SVD | TOBMI | Lasso | TDimpute-self | TDimpute | SVD | TOBMI | Lasso | TDimpute-self | TDimpute |
| 10% | 65 | 64.2 | 63 | 67.8 | **68** | 59.8 | 63 | 63 | 63.4 | **68.8** |
| 30% | 45.2 | 48 | 48.2 | **53.2** | 50.2 | 32.6 | 35.4 | 34.8 | **38.6** | **38.6** |
| 50% | 29.8 | 32.2 | 33.4 | **37** | 34.2 | 20.4 | 20.2 | 20.2 | 23.6 | **24.8** |
| 70% | **23.2** | 19 | 21.4 | 21.4 | 22.2 | 10.4 | 9.4 | 10 | 11 | **14.2** |
| 90% | 7.2 | 4.6 | 5.8 | 8 | **13** | 2.2 | 2 | 2.8 | 2.8 | **6** |
| Missing rate | CESC |  |  |  |  | KIRP |  |  |  |  |
|  | SVD | TOBMI | Lasso | TDimpute-self | TDimpute | SVD | TOBMI | Lasso | TDimpute-self | TDimpute |
| 10% | 81.6 | 79.6 | 82.8 | 80.2 | **82.2** | 88.6 | 87.8 | 77.4 | 87.8 | **88.8** |
| 30% | 58.8 | 60.4 | 59.2 | 59 | **63.6** | **85.2** | 68 | 53.2 | 85 | 81 |
| 50% | 43.6 | 47.2 | 47 | 48.4 | **50.4** | **83.8** | 55.6 | 36.8 | 82.2 | 77.6 |
| 70% | 21.2 | 21.4 | 26 | **31.2** | 30.2 | 80.2 | 57.4 | 30.6 | **83.2** | 82 |
| 90% | 4.4 | 5.4 | 5.4 | 8.4 | **13** | 49.4 | 49.6 | 26.8 | 59.4 | **67.4** |
| Missing rate | LGG |  |  |  |  | HNSC |  |  |  |  |
|  | SVD | TOBMI | Lasso | TDimpute-self | TDimpute | SVD | TOBMI | Lasso | TDimpute-self | TDimpute |
| 10% | 90.2 | 89.6 | 76.4 | 91.2 | **92.6** | 69 | 70.6 | 71.8 | **77.2** | **77.2** |
| 30% | 87.4 | 87.2 | 66.4 | **88.6** | 88 | 47 | 50.4 | 48.4 | 52.2 | **56** |
| 50% | 88 | 86.4 | 55.2 | **89** | 88 | 28.2 | 33.6 | 31.2 | 37.6 | **42.4** |
| 70% | 84.4 | 90.4 | 61.4 | 90.6 | **91.2** | 16.4 | 18.8 | 18 | 22.2 | **28.4** |
| 90% | 80.8 | 85.4 | 65.2 | **92.6** | 90 | 6.4 | 3.6 | 4.4 | 7.2 | **13.8** |
| Missing rate | SKCM |  |  |  |  | BLCA |  |  |  |  |
|  | SVD | TOBMI | Lasso | TDimpute-self | TDimpute | SVD | TOBMI | Lasso | TDimpute-self | TDimpute |
| 10% | 61 | 66.2 | 0 | **66.4** | 65.2 | 69.4 | 74.6 | 74.2 | 74.8 | **75.6** |
| 30% | 30.2 | 37.2 | 0.8 | 37.4 | **39.2** | 37.6 | 47.6 | 49.4 | **54.6** | 54.2 |
| 50% | 11.8 | 15.2 | 0.6 | **21** | 20.4 | 21.2 | 26.8 | 31 | **41.8** | 41.6 |
| 70% | 3.2 | 6.2 | 0 | 8.6 | **11.8** | 14.6 | 15.2 | 18 | 27.2 | **31.8** |
| 90% | 1.8 | 0.4 | 0 | 1.4 | **4.4** | 7.8 | 4.6 | 7.6 | 10.2 | **20.6** |
| Missing rate | STAD |  |  |  |  | KIRC |  |  |  |  |
|  | SVD | TOBMI | Lasso | TDimpute-self | TDimpute | SVD | TOBMI | Lasso | TDimpute-self | TDimpute |
| 10% | 71.6 | 70.2 | 69 | 72.2 | **73** | 91.4 | 89.6 | 84.6 | **91.4** | 91.2 |
| 30% | 37.8 | 42.4 | 42.2 | 44.4 | **45.2** | 75.8 | 72.6 | 55 | 81.8 | **83.2** |
| 50% | 25.4 | 26.4 | 25.2 | 25.8 | **29.6** | 65.4 | 64.4 | 36.6 | **81.2** | 78.2 |
| 70% | 8.2 | 9.4 | 7.2 | 12.2 | **12.6** | 57.8 | 58.6 | 32.8 | **76.6** | 72.4 |
| 90% | **3.8** | 3.4 | 1.8 | 3.2 | 3.6 | 38.6 | 49.8 | 24.4 | **63.6** | 62 |
| Missing rate | COAD |  |  |  |  | SARC |  |  |  |  |
|  | SVD | TOBMI | Lasso | TDimpute-self | TDimpute | SVD | TOBMI | Lasso | TDimpute-self | TDimpute |
| 10% | 59.8 | 63.4 | 62.2 | 64.8 | **65.6** | 73.4 | 77 | 75.8 | 79.4 | **79.6** |
| 30% | 30.4 | 33 | 30.8 | 32.8 | **37.8** | 45.8 | 57.2 | 53.8 | 57.8 | **58** |
| 50% | 18.4 | **21** | 20.4 | 19.2 | 20.6 | 30 | 43 | 37.8 | 46 | **49** |
| 70% | 7.8 | 6.6 | 8.4 | 9.2 | **15** | 20.8 | 30 | 29 | 35.2 | **37.2** |
| 90% | 2 | 2.4 | 1.4 | 2.4 | **8.2** | 3.2 | 18.2 | 10.4 | 23.2 | **27** |

The results are averaged over 5 random replicas. Best results are highlighted in bold face.

**Table S5.** The enrichment factors of the top 100 ranked genes in the gene list from The Human Protein Atlas across 16 cancer types

| Missing rate | BRCA |  |  |  |  | THCA |  |  |  |  |
| --- | --- | --- | --- | --- | --- | --- | --- | --- | --- | --- |
|  | SVD | TOBMI | Lasso | TDimpute-self | TDimpute | SVD | TOBMI | Lasso | TDimpute-self | TDimpute |
| 10% | 7.637 | 8.625 | 8.098 | 8.493 | **8.888** | 8.664 | 9.102 | **9.322** | 8.554 | 9.102 |
| 30% | 4.411 | **6.584** | 5.925 | 6.452 | 6.386 | 5.045 | 6.251 | 6.141 | **6.799** | 6.361 |
| 50% | 2.765 | 4.740 | 3.489 | **5.267** | **5.267** | 3.400 | 4.058 | 4.167 | **4.496** | **4.496** |
| 70% | 1.383 | 2.436 | 1.185 | 3.226 | **3.424** | 2.303 | 2.961 | 2.632 | **3.509** | 2.522 |
| 90% | 0.790 | 0.988 | 0.593 | 1.580 | **1.843** | 1.755 | 1.206 | 0.987 | **2.303** | 1.864 |
| Missing rate | PRAD |  |  |  |  | LUAD |  |  |  |  |
|  | SVD | TOBMI | Lasso | TDimpute-self | TDimpute | SVD | TOBMI | Lasso | TDimpute-self | TDimpute |
| 10% | 14.60 | **15.08** | **15.08** | 13.40 | 14.12 | 8.476 | 8.476 | 8.242 | **8.944** | **8.944** |
| 30% | 8.86 | **9.57** | 9.33 | 8.86 | 8.14 | 6.839 | 6.079 | 5.962 | **7.541** | **7.541** |
| 50% | 4.31 | **5.74** | 5.50 | 4.55 | 4.55 | 5.436 | 4.618 | 4.559 | 5.904 | **5.962** |
| 70% | 3.11 | **4.79** | 3.35 | 3.59 | 2.15 | 3.566 | 2.923 | 3.040 | 4.150 | **5.904** |
| 90% | 0.96 | 1.68 | 1.44 | **2.15** | 1.44 | 2.455 | 0.877 | 1.695 | 2.747 | **4.092** |
| Missing rate | LIHC |  |  |  |  | LUSC |  |  |  |  |
|  | SVD | TOBMI | Lasso | TDimpute-self | TDimpute | SVD | TOBMI | Lasso | TDimpute-self | TDimpute |
| 10% | 1.004 | 0.951 | 0.977 | 1.017 | **1.030** | 1.403 | 1.637 | 1.461 | 1.578 | **1.695** |
| 30% | **0.700** | 0.647 | 0.687 | **0.700** | 0.674 | 0.760 | 1.169 | 0.877 | 1.286 | **1.461** |
| 50% | 0.396 | 0.449 | **0.489** | 0.436 | 0.370 | 0.526 | 0.760 | 0.468 | 0.701 | **0.818** |
| 70% | 0.304 | 0.225 | **0.330** | 0.225 | 0.277 | 0.292 | **0.526** | 0.234 | 0.292 | 0.468 |
| 90% | **0.119** | 0.000 | 0.066 | 0.092 | **0.119** | 0.000 | **0.058** | 0.000 | 0.000 | **0.058** |
| Missing rate | CESC |  |  |  |  | KIRP |  |  |  |  |
|  | SVD | TOBMI | Lasso | TDimpute-self | TDimpute | SVD | TOBMI | Lasso | TDimpute-self | TDimpute |
| 10% | 13.09 | 13.19 | 13.30 | 13.19 | **13.51** | 0.363 | 0.363 | 0.300 | **0.370** | 0.363 |
| 30% | 9.62 | 10.62 | 9.88 | 10.04 | **10.83** | **0.351** | 0.287 | 0.223 | **0.351** | 0.332 |
| 50% | 6.73 | 8.15 | 7.78 | 7.99 | **8.83** | **0.370** | 0.236 | 0.134 | **0.370** | 0.338 |
| 70% | 2.58 | 3.68 | 3.94 | 4.73 | **5.10** | 0.332 | 0.300 | 0.147 | **0.363** | 0.338 |
| 90% | 0.47 | 0.68 | 0.95 | 1.31 | **2.05** | 0.217 | 0.217 | 0.108 | 0.255 | **0.281** |
| Missing rate | LGG |  |  |  |  | HNSC |  |  |  |  |
|  | SVD | TOBMI | Lasso | TDimpute-self | TDimpute | SVD | TOBMI | Lasso | TDimpute-self | TDimpute |
| 10% | NA |  |  |  |  | 6.29 | 6.92 | 6.97 | **7.45** | 7.40 |
| 30% |  |  |  |  |  | 3.84 | 4.71 | 4.71 | 5.00 | **5.53** |
| 50% |  |  |  |  |  | 1.97 | 3.32 | 2.64 | 3.80 | **4.23** |
| 70% |  |  |  |  |  | 0.96 | 1.63 | 1.35 | 2.21 | **2.64** |
| 90% |  |  |  |  |  | 0.34 | 0.19 | 0.24 | 0.72 | **1.01** |
| Missing rate | SKCM |  |  |  |  | BLCA |  |  |  |  |
|  | SVD | TOBMI | Lasso | TDimpute-self | TDimpute | SVD | TOBMI | Lasso | TDimpute-self | TDimpute |
| 10% | 2.78 | **3.53** | 0 | 2.78 | 3.16 | 6.65 | 7.24 | 7.45 | 7.28 | **7.56** |
| 30% | 1.11 | **2.41** | 0 | 2.23 | 1.86 | 3.13 | 4.94 | 4.91 | **5.74** | 5.71 |
| 50% | 0.19 | 1.11 | 0 | 0.93 | **1.49** | 1.67 | 2.82 | 2.85 | **4.42** | 4.14 |
| 70% | 0.00 | 0.74 | 0 | 0.00 | **1.11** | 0.91 | 1.57 | 1.85 | 2.79 | **3.31** |
| 90% | 0.00 | 0.00 | 0 | 0.00 | **0.37** | 0.59 | 0.35 | 0.59 | 0.97 | **2.26** |
| Missing rate | STAD |  |  |  |  | KIRC |  |  |  |  |
|  | SVD | TOBMI | Lasso | TDimpute-self | TDimpute | SVD | TOBMI | Lasso | TDimpute-self | TDimpute |
| 10% | 5.09 | 5.09 | 5.09 | **5.47** | 5.35 | 0.281 | 0.281 | 0.268 | **0.287** | **0.287** |
| 30% | 2.80 | 2.80 | 2.67 | 3.18 | **3.69** | 0.255 | 0.261 | 0.166 | 0.255 | **0.268** |
| 50% | 0.89 | 1.65 | 1.40 | 1.53 | **1.78** | 0.210 | 0.223 | 0.077 | 0.268 | **0.274** |
| 70% | 0.00 | 0.38 | 0.51 | **0.89** | 0.76 | 0.185 | 0.204 | 0.089 | 0.249 | **0.268** |
| 90% | 0.00 | 0.13 | 0.13 | 0.00 | **0.25** | 0.096 | 0.185 | 0.038 | 0.217 | **0.236** |
| Missing rate | COAD |  |  |  |  | SARC |  |  |  |  |
|  | SVD | TOBMI | Lasso | TDimpute-self | TDimpute | SVD | TOBMI | Lasso | TDimpute-self | TDimpute |
| 10% | 1.02 | 1.09 | 1.09 | 1.15 | **1.28** | NA |  |  |  |  |
| 30% | 0.77 | **0.83** | **0.51** | 0.70 | 0.77 |  |  |  |  |  |
| 50% | 0.32 | 0.45 | 0.26 | **0.51** | 0.32 |  |  |  |  |  |
| 70% | 0.00 | 0.00 | 0.00 | 0.00 | **0.19** |  |  |  |  |  |
| 90% | 0.06 | 0.06 | 0.00 | 0.06 | **0.13** |  |  |  |  |  |

The results are averaged over 5 random replicas. Best results are highlighted in bold face.
LGG and SARC are not included in this scenario, because no relevant gene list provided by The Human Protein Atlas.

**Fig S7.**
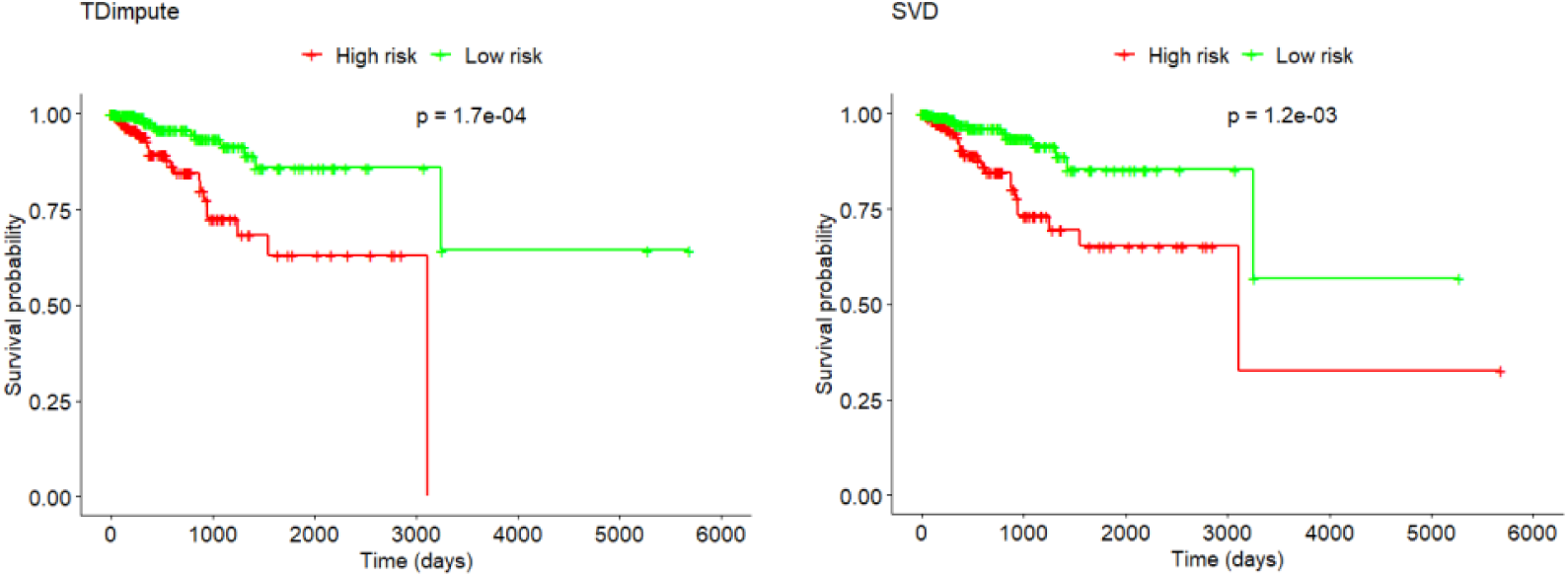
Kaplan-Meier plot for the two clusters obtained from the UCEC dataset imputed by TDimpute and SVD, respectively.

**Fig S8.**
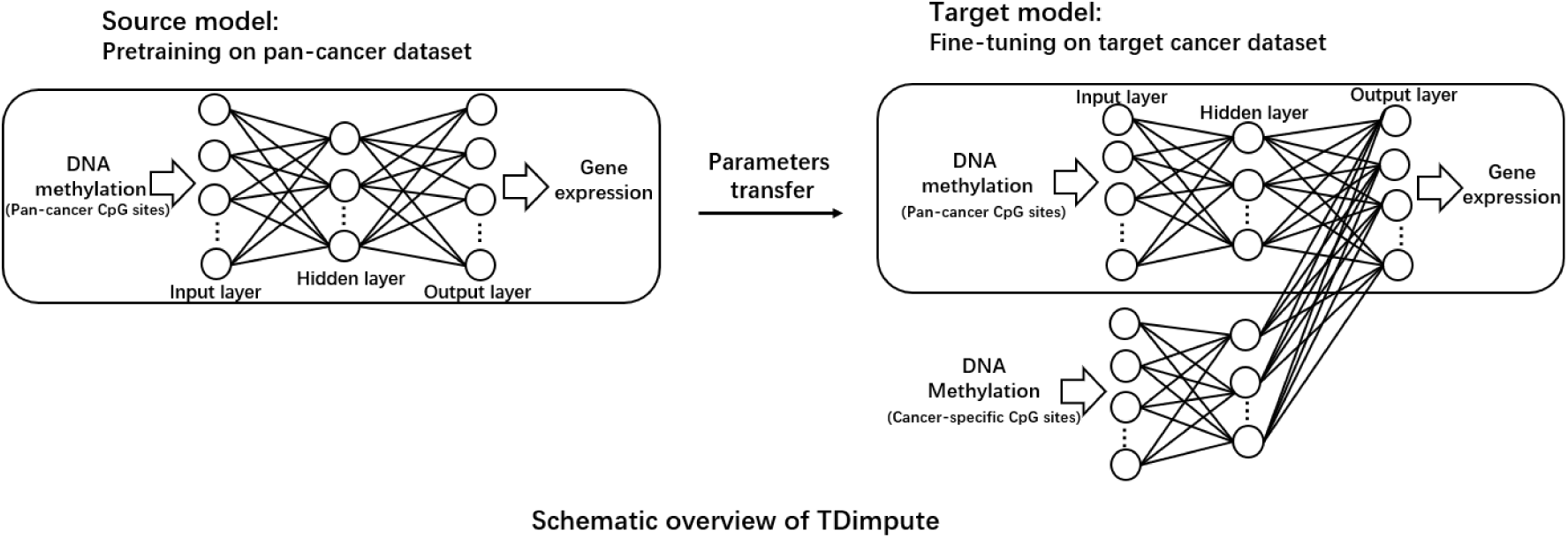
The architecture of transfer learning based neural network (TDimpute) with cancer-specific CpG sites as auxiliary input. The input at fine-tuning stage consists of two parts: the cancer-specific part takes the highly variable CpG sites from the target cancer as input, and the transfer part takes the commonly variable CpG sites from pan-cancer dataset as input.

**FigS9.**
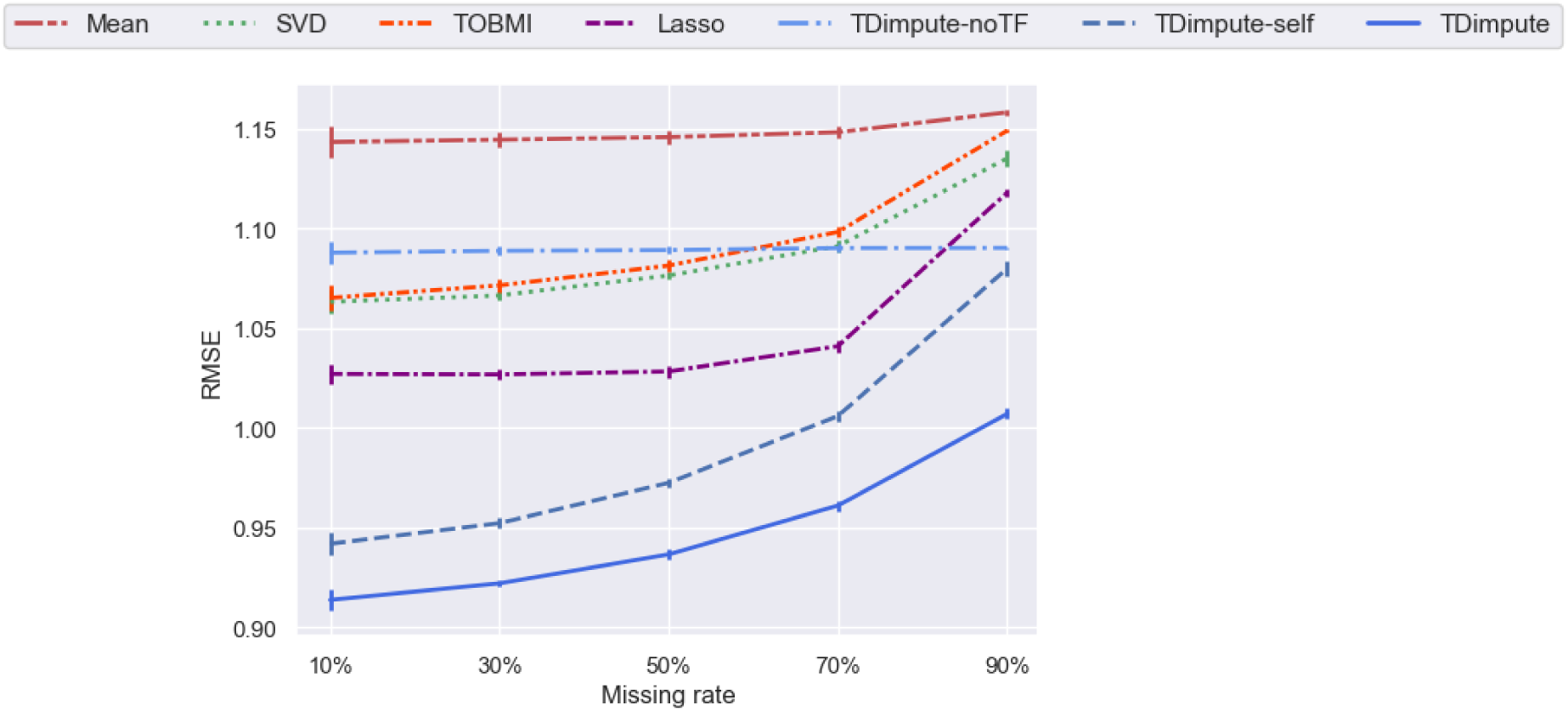
RMSE values of each imputation method with top 20000 CpG sites as input. For SVD, TOBMI, Lasso and TDimpute-self, the top 20000 variable cancer-specific CpG sites are used as input, while the input of TDimpute includes both the pan-cancer and cancer-specific CpG sites. Results were averaged across 16 imputed cancer datasets. The error bar shows the standard deviation.

**Fig S10.**
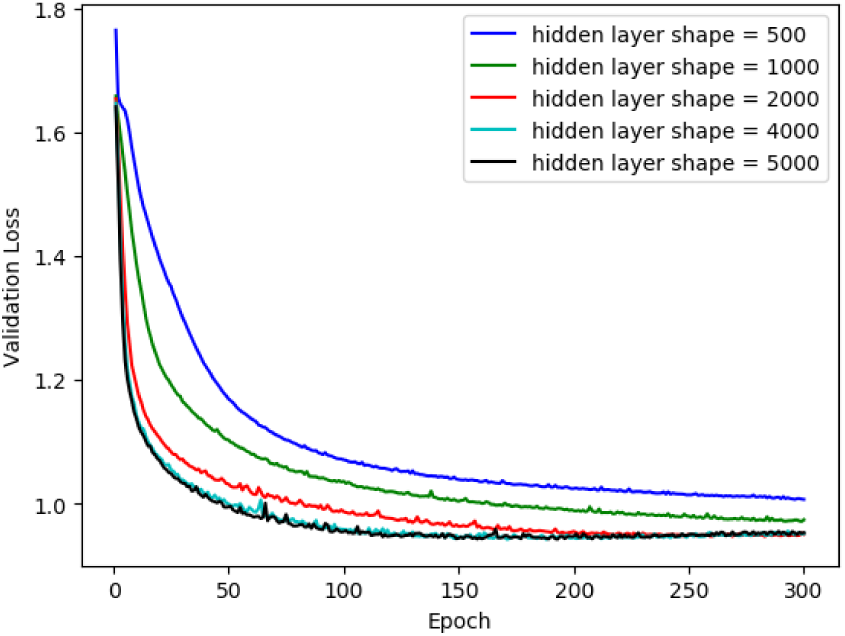
Loss curves for different hidden layer shape

**Table S6.** hyper-parameter analysis for hidden layer shape, the number of hidden layers, and training epochs on pan-cancer dataset (excluding BRCA dataset).

|  |  | epoch |  |  |  |  |
| --- | --- | --- | --- | --- | --- | --- |
| the number of hidden layers | hidden layer shape | 50 | 100 | 150 | **300** | 500 |
| **1** | 500 | 1.253 | 1.131 | 1.09 | 1.052 | 1.04 |
|  | 1000 | 1.146 | 1.076 | 1.047 | 1.016 | 1.013 |
|  | 2000 | 1.074 | 1.029 | 1.009 | 0.995 | 1.009 |
|  | **4000** | 1.043 | 1.003 | 0.991 | 0.997 | 1.015 |
|  | 5000 | 1.035 | 0.999 | 0.988 | 0.995 | 1.015 |
| 3 | (4000, 500, 4000) | 1.13 | 1.067 | 1.039 | 1.01 | 1.011 |
|  | (4000, 1000, 4000) | 1.107 | 1.046 | 1.032 | 1.011 | 1.035 |
|  | (4000, 2000, 4000) | 1.092 | 1.042 | 1.021 | 1.025 | 1.064 |
|  | (5000, 500, 5000) | 1.085 | 1.034 | 1.021 | 1.028 | 1.078 |
|  | (5000, 1000, 5000) | 1.097 | 1.052 | 1.025 | 1.014 | 1.04 |
|  | (5000, 2000, 5000) | 1.124 | 1.062 | 1.036 | 1.008 | 1.012 |

Selected hyper-parameters are highlighted in bold face.

**Fig S11.**
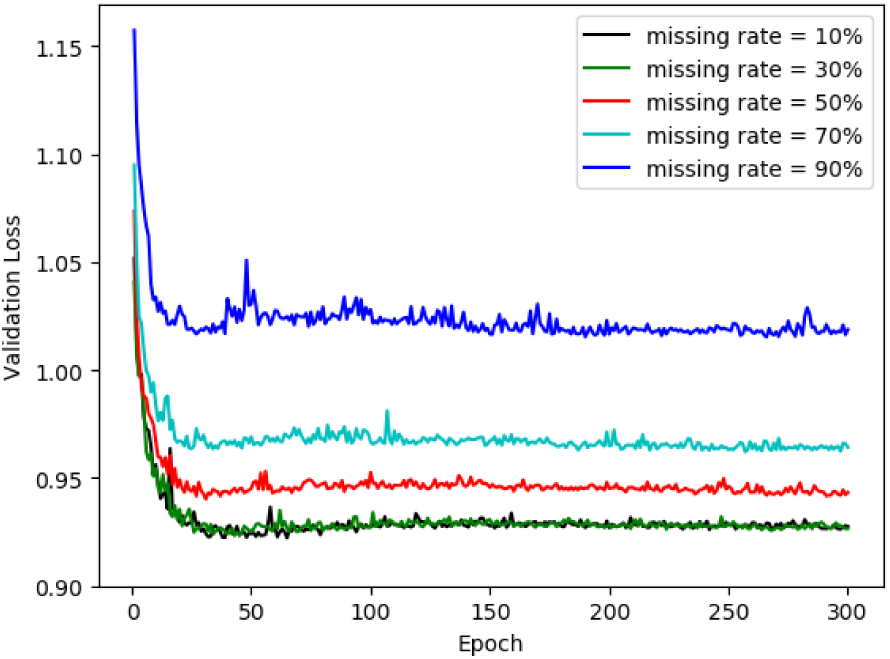
The loss curves of different missing rates on the validation dataset of BRCA.

